# Noise-Reducing Negative-Feedback Optogenetic Circuits in Mammalian Cells

**DOI:** 10.1101/601005

**Authors:** Michael Tyler Guinn, Gábor Balázsi

## Abstract

Gene autorepression is widely present in nature and is also employed in synthetic biology, partly to reduce gene expression noise in cells. Optogenetic systems have recently been employed for controlling gene expression levels in mammalian cells, but most have utilized activator-based proteins, neglecting negative feedback. Here, we engineer optogenetic negative-feedback gene circuits in mammalian cells to achieve noise-reduction for precise gene expression control. We build a toolset of these noise-reducing Light-Inducible Tuner (LITer) gene circuits using the TetR repressor fused with a Tet-Inhibitory peptide (TIP) or a degradation tag through the light-sensitive LOV2 protein domain. These LITers provide nearly a range of 4-fold gene expression control and up to five-fold noise reduction from existing optogenetic systems. Moreover, we use the LITer gene circuit architecture to control gene expression of the cancer oncogene KRAS(G12V) and study its downstream effects through phospho-ERK levels and cellular proliferation. Overall, these novel LITer optogenetic platforms should enable precise spatiotemporal perturbations for studying multicellular phenotypes in developmental biology, oncology, and other biomedical fields of research.

## Main

Gene expression levels and variability (noise) dictate transcript and protein production which define the properties of living cells in health and disease^1,2^. Depending on the interplay between gene function and environmental conditions, expression levels and noise in cellular populations can confer a variety of cellular advantages and disadvantages^3–8^. The importance of gene expression noise in cellular processes may be one reason why natural gene regulatory networks contain noise-modulating network motifs^9–11^. For example, Negative-Feedback (NF) is a critical and frequent network motif that can reduce noise^11–14^ in biological processes as diverse as circadian rhythm, immunological responses, or stress signaling in cancer^15–17^. Engineering gene circuits that control gene expression levels and noise simultaneously can reveal important thresholds and sensitivities for broad biological phenomena such as metastasis, epithelial-to-mesenchymal (EMT) transition, and drug resistance^18^.

A few decades ago, bacterial regulator-based systems emerged capable of controlling intermediate gene expression levels^19–21^. Despite this advancement, these systems often suffered from high noise since they lacked feedback regulation, therefore leaving gene expression variability as an uncontrolled parameter^22^. Adjusting noise has also been neglected by traditional methods of gene expression control, which tend to focus on extreme gene expression changes (e.g. knockout and overexpression) in cells or organisms.

Engineered solutions have emerged more recently using NF in synthetic gene circuits to fine-tune protein expression proportional to an extracellular chemical inducer while also reducing noise^23^. However, existing NF gene circuits respond relatively slowly to chemical stimulus and do not enable single-cell level gene expression control^22–25^. Optogenetic systems have the potential to overcome these limitations within living cells^26,27^. Yet, most optogenetic gene circuit studies have relied on transient transfection and their ability to deliver spatiotemporal control with low noise in single cells remains unknown^28,29^. Additionally, most existing optogenetic systems are activator-based, incompatible with noise-reduction by NF^28,29^.

Here, as a step towards precisely controlling single-cells with light, we engineer a series of stable human cell lines expressing NF optogenetic gene circuits developed from previously constructed ‘Linearizers’^23,28^. To achieve this, we introduce two novel peptide elements for controlling the Tet-repressor (TetR). We fuse TetR with a light-responsive protein domain (LOV2)^27,30,31^, which is further fused to either a Tet-inhibiting peptide (TIP)^32^ or a degradation tag consisting of four amino acids^33^. These components remain hidden until illuminated, at which point they ensure light-dependent inhibition or degradation of the TetR protein^32–34^. The modified TetR regulators repress a stably-integrated fluorescent reporter, enabling light-dependent gene expression measurements. To assess the performance of these novel optogenetic systems, we compare them with the existing optogenetic LightOn system^28^ integrated into the same parental cell line. Lastly, we also study the versatility of these optogenetic circuits for controlling the mutated KRAS(G12V) gene, relevant to pancreatic and colon cancer^35,36^.

Overall, we present novel optogenetic tools that can enable a wide-dynamic range of gene expression in response to light stimuli and provide low noise for controlling single mammalian cells. These synthetic biology tools have the potential to be utilized in single-cell studies of processes as diverse as embryonic development, cancer metastasis, and neuron migration.

## Results

### Testing the precision of gene expression control with an established optogenetic gene circuit

To study the precision of gene expression control for an existing optogenetic benchmark system, we first constructed a stable Flp-In™ 293 cell line integrating the LightOn system expressing the green fluorescence protein, mNeonGreen (Figure 1A & Supporting Figure S1A)^28^. Previous studies have explored gene expression of the LightOn system with transient transfections but have not analyzed stably-integrated mammalian cell-lines^37–39^. Subsequently, we will refer to this stable engineered cell line as VVD.

**Figure 1.**
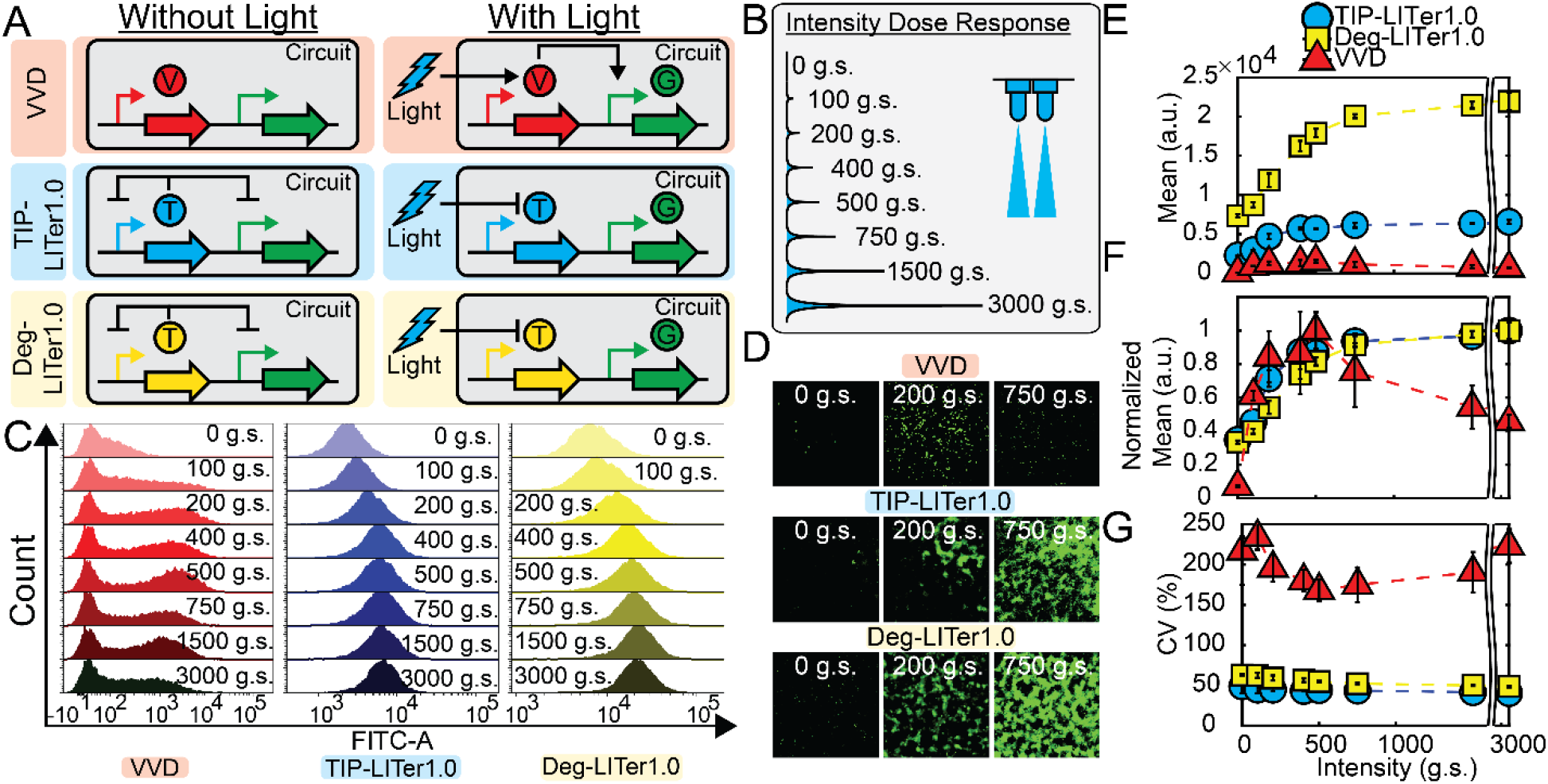
Gene expression control by VVD, TIP-LITer1.0, & Deg-LITer1.0 gene circuits with constant illumination at increasing intensities. (A) Schematic illustration of VVD, TIP-LITer1.0, & Deg-LITer1.0 gene circuits stably integrated within Flp-In 293 cell genome. ‘V’ is the VVD protein, ‘T’ is the TetR regulator with either TIP or Degron, and ‘G’ is the Reporter output. (B) Schematic illustration of experimental design with cells exposed to varying blue light intensities. (C) Fluorescence histogram distributions for flow cytometry performed for each corresponding gene circuit under light intensity titration. (D) Fluorescence microscopy of cells with corresponding gene circuits over light intensity titration. (E) Flow cytometry mean fluorescence expression versus light intensity titration for the three gene circuits. (F) Normalized mean fluorescence values to max light intensity level for each gene circuit. Error bars are standard deviation, N=3. (G) Coefficient of variation (CV) over light intensity titration for the three gene circuits. Two-sample t-test was performed on maximum and minimum mean fluorescence data values. TIP-LITer1.0 ON/OFF had a p-value of 3.21E-06, Deg-LITer1.0 ON/OFF had a p-value of 5.40E-10, and VVD ON/OFF had a p-value of 1.28E-04. Kruskal-Wallis test was performed on CV data set differences. TIP-LITer1.0 vs VVD had a p-value of 2.87E-09 and Deg-LITer1.0 vs VVD had a p-value of 2.88E-09.

To characterize the light responsiveness of VVD cells, we sought a robust illumination platform that could allow a wide-range of tunability for various light parameters. Thus, we turned to the recent Light Plate Apparatus (LPA) system, which is easily customizable for *in vitro* assays, allowing parameter scans for light intensities, pulses, and complex light patterns^40^. We constructed several LPA systems allowing up to 24-light conditions to be probed simultaneously per device (Supporting Figure S2).

To explore VVD cell response to light, we first analyzed gene expression under increasing light intensities with continuous illumination (Figure 1B). We observed nearly 14-fold change of gene expression induction (Figure 1C). However, around the LPA induction of 500 g.s., the response of VVD cells reached a maximum and then dropped for increasing light intensities (Supporting Figure S3). This behavior has not been described before, potentially because previous studies have utilized custom light apparatuses with limited light probing capabilities. Moreover, we also noticed large gene expression noise, with a coefficient of variation (CV) approximately 5-fold higher compared to existing chemically-inducible NF gene circuits^23^. This observation led us to create light-controllable systems with low noise since high CV may be non-optimal for precise single-cell gene expression control.

### LITer1.0 gene circuits improve the precision of single-cell gene expression control

To achieve lower noise than observed in the VVD cells, we began exploring gene circuit designs that reduce noise. Considering the noise-reducing ability of NF, we moved to integrate negative autoregulation into optogenetic circuits. Notably, most optogenetic systems currently rely on transcriptional activation, which is incompatible with negative autoregulation^28,29^.Thus, we turned to existing chemical-inducible mammalian gene expression systems that utilized TetR negative feedback to precisely control GFP reporter expression^23^. We hypothesized that upgrading these chemical tools into optogenetic systems could replicate their wide dynamic range of gene expression induction with low noise^23^.

To enable NF in optogenetics, we turned towards the blue-light responsive (450 nm) LOV2 protein domain which has been fused to various proteins previously^27,31^. We fused the LOV2 protein domain with the repressor protein TetR (Figure 1A) and added the Tet-inhibiting peptide (TIP)^32^ capable of inhibiting the TetR protein to the C-terminal end of LOV2. We reasoned that the LOV2 protein domain should open-up upon blue light stimulation, exposing the TIP to bind and inactivate TetR, de-repressing GFP and enabling its expression. Finally, we incorporated this TetR-LOV2-TIP fusion protein under the control of a CMV-based D2ir promoter with two Tet operator (*TetOx2*) sites, allowing feedback regulation^23^. Additionally, an identical D2ir promoter drove the expression of the reporter GFP (Figure 1A & Supporting Figure S1B). We constructed the entire gene circuit into a single vector, stably integrated into Flp-In™ 293 cells, and sorted for monoclonal cells. Henceforth, we refer to the cells expressing this gene circuit as TIP-based Light-Inducible Tuner or TIP-LITer1.0.

To complement the TIP-LITer1.0, we next modified the gene circuit by replacing the TIP with a degradation tag consisting of four amino acids (RRRG, Figure 1A & Supporting Figure S1C)^33^. Like the TIP-LITer1.0 system, the gene circuit is within the OFF-state under darkness. Upon blue light stimulation, the LOV2 protein domain opens and exposes the RRRG tag leading to degradation of the TetR fusion repressor, allowing the expression of GFP. Henceforth, the cells expressing this gene circuit will be referred to as Degron-based Light-Inducible Tuner, or Deg-LITer1.0.

Next, to compare the characteristics of these new gene circuits with an existing optogenetic circuit, we performed the same light intensity dose-response measurements on TIP-LITer1.0 and Deg-LITer1.0 as we did with the VVD system. We observed a wide dynamic range of gene expression induction (Figure 1C & D). Importantly, at the population level, we observed much tighter fluorescence distributions for both the TIP-LITer1.0 & the Deg-LITer1.0, indicating lower gene expression noise.

For quantitative comparisons of the three gene circuits’ gene expression levels, we examined the mean fluorescence output (Figure 1E & F). Interestingly, the Deg-LITer1.0 had the highest expression, the TIP-LITer1.0 had an intermediate expression, and the VVD had the lowest level of expression. We achieved approximately 3-fold dynamic gene expression induction for all light doses for both LITer1.0 systems as opposed to the VVD system, which became inactivated at prolonged periods of high-intensity light exposure (Supporting Figure S3). Consequently, we performed measurements on the VVD system at 24-hour terminal time-points and the LITer1.0 systems at 72-hour terminal time points.

To compare the noise of the three systems we examined their CV. Strikingly, we observed up to 5.5 and 4.5-fold lower CV of TIP-LITer1.0 & Deg-LITer1.0 compared with the VVD system (Figure 1G). We hypothesize this noise reduction may have three sources. First, negative feedback is known to reduce gene expression noise^12,41^. Second, the slow kinetics of the VVD protein versus the fast kinetics of the LOV2 protein domain may lead to higher variation among induced VVD cells^42,43^. Third, monomers of the LOV2 protein domain can induce gene expression while VVD activation requires dimerization, possibly increasing cooperativity and noise amplification.

To explore the use of LITer1.0 gene circuits as analog tools for controlling gene expression, we also inspected the linearity of light intensity dose-response (Supporting Information) ^41^. The LITer1.0 dose-responses were approximately linear up to 225 g.s. from the LPA (Supporting Figure S4), complementing other systems that exhibit linearity with pulsed illumination^44^. The linearity was somewhat reduced in the LITer1.0 gene circuits compared to chemically-inducible systems, because of two main dose-response features. First, the gene circuit did not respond at very low light intensities, causing the dose-response to start off flat (Supporting Figure 4). Second, even after this flat part, the dose-response did not remain linear for a wide inducer range, starting to curve downwards at ~20% (75 g.s.) saturation (Supporting Figure 4). To investigate the sources of these deviations from standard linearizer characteristics, we generalized earlier deterministic models^23^, allowing basal gene expression, reversion of induced LOV2 into its inactive state, and adjustment of the Hill parameters. Indeed, elevated basal gene expression explained the flat start-off, while Hill parameter adjustments captured the reduced linearity (Supporting Figure 5 & Supporting information). Additionally, increased refolding of LOV2 into an inactive state increased the light needed to induce a specific level of GFP. When combining all these parameters into a single model (Supporting Figure 5), the dose response more closely matched the experimental results.

Overall, these intensity dose-response experiments provide initial characterization of a classical and two newly engineered optogenetic systems, which we next explored under pulsed light-induction regimes.

### Responses of VVD and LITer1.0 gene circuits to pulsed inputs

To characterize the response of VVD and LITer1.0 gene circuits to discontinuous light-induction regimes, we next investigated by fluorescence microscopy and flow cytometry measurements how single light pulses of variable duration and fixed LPA intensity (1000 g.s.) affects gene expression and noise (Figure 2A & Supporting Figure S6).

**Figure 2.**
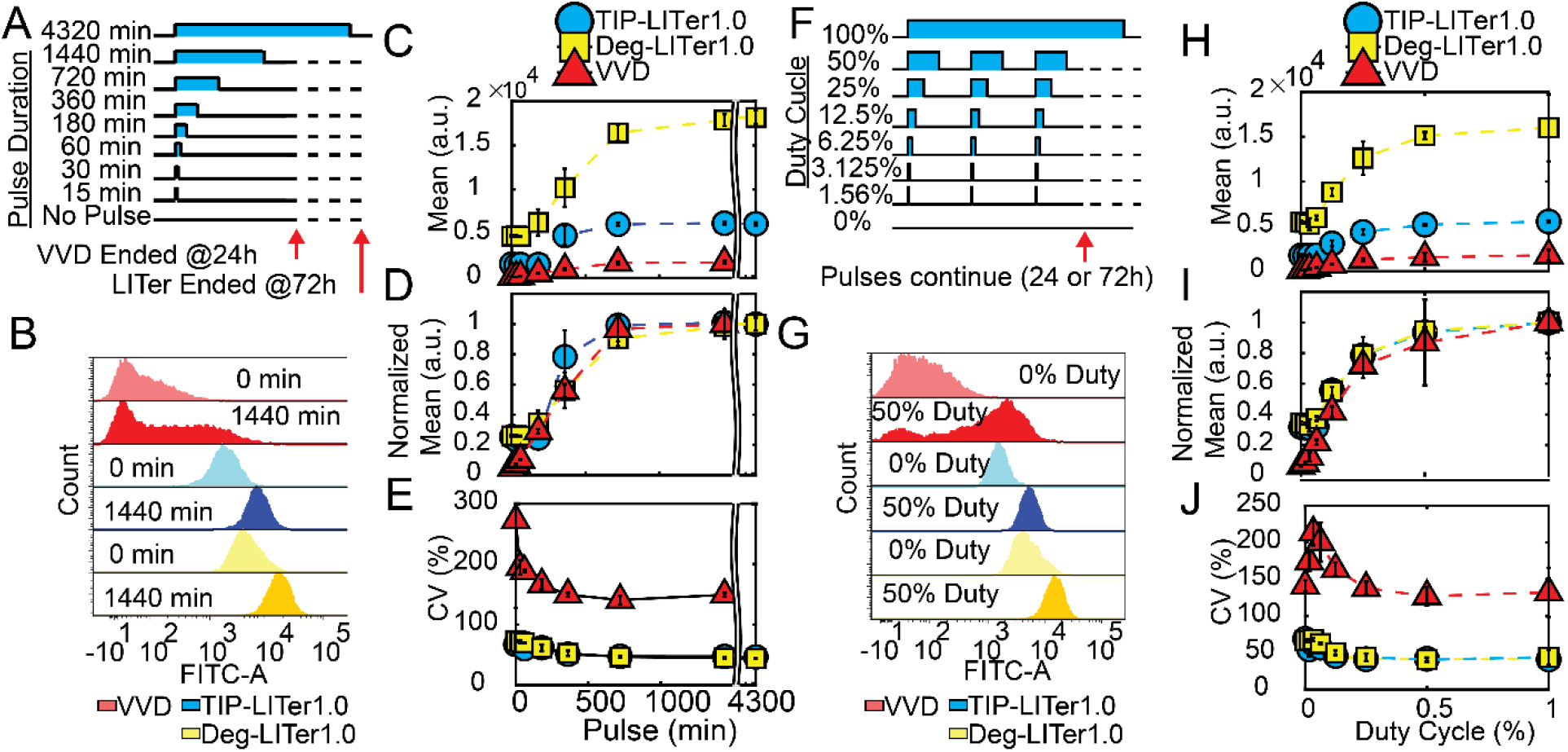
Gene expression control by VVD, TIP-LITer1.0, & Deg-LITer1.0 gene circuits with pulsed illumination. (A) Schematic illustration of single pulse experiment. (B) Fluorescence histogram distributions for single light pulse titration. (C) Flow cytometry mean fluorescence expression for single pulse experiment. (D) Normalized mean fluorescence values for single pulse experiment. (E) Coefficient of variation (CV) for single pulse experiment. (F) Schematic illustration of duty cycle experiment. (G) Fluorescence histogram distributions for duty cycle experiment. (H) Flow cytometry mean fluorescence for duty cycle experiment. (I) Normalized mean fluorescence values for duty cycle experiment (J) CV over duty cycle experiment. Error bars are standard deviation, N=3. Two-sample t-test was performed on maximum and minimum mean fluorescence data values for pulse and duty-cycle experiments. TIP-LITer1.0 ON/OFF had a p-value of 1.76E-06, Deg-LITer1.0 ON/OFF had a p-value of 7.77E-06, and VVD ON/OFF had a p-value of 7.03E-05 for pulse experiment. TIP-LITer1.0 ON/OFF had a p-value of 3.17E-06, Deg-LITer1.0 ON/OFF had a p-value of 0.0083, and VVD ON/OFF had a p-value of 0.0092 for duty-cycle experiment. Kruskal-Wallis test was performed on CV data set differences. TIP-LITer1.0 vs VVD and Deg-LITer1.0 vs VVD had a p-value of 2.88E-09 for the pulse experiment. TIP-LITer1.0 vs VVD had a p-value of 2.88E-09 and Deg-LITer1.0 vs VVD had a p-value of 2.87E-09 for the duty-cycle experiment.

We found nearly 4-fold change of gene expression induction for the LITer1.0 gene circuits compared to 21.9 for VVD for different pulse lengths. Interestingly, VVD expression saturated for pulsed illumination (Figure 2B-D), implying that shorter (24 hour) illuminations and intermediate intensities eliminate the fall in activity at high doses with continuous illumination. Importantly, we also observed around 4-fold lower gene expression noise for both LITer1.0 gene circuits compared with the VVD system (Figure 2E).

As another discontinuous light-induction regime, we tested the effects of varying the duty cycle (percent of time ON) for periodic light stimuli on the gene circuits. We chose a stimulus period of 1 hour and measured gene expression at increasing duty cycle percentages (Figure 2F). Again, we found broad distributions in the VVD system and tight distributions for the LITer1.0 systems (Figure 2G). The VVD system could achieve around 2-fold induction with as little as 6% duty cycle for which the LITer1.0 systems required 12.5% duty cycle (Figure 2H & I). As before, the CV of the LITer1.0 gene circuits were several-fold lower (Figure 2J).

Interestingly, low duty cycles could achieve near maximal expression implying there may be benefits to low ON/OFF ratios versus continuous light, perhaps due to cellular toxicity at long exposures. Overall, the LITer1.0 gene circuits had lower noise than VVD, but their basal expression was high for expressing functional genes. To optimize these optogenetic circuit prototypes for functional gene expression, we next turned to modeling to reveal experimental parameters that could be adjusted to lower basal expression.

### Computational Models suggest improvements for LITer1.0 gene circuits

To examine strategies for optimizing the LITer1.0 architecture by lowering basal expression, we asked if reducing the system from two promoters to one promoter would lower the basal expression. We reasoned that decreasing the number of *TetO* operator sites competing for TetR might increase the effective time TetR would be bound and lead to stronger repression of GFP expression. Therefore, we developed deterministic and stochastic models of basal expression in double-promoter LITer1.0 systems versus single-promoter LITer2.0 systems, considering a set of simple reactions including transcription, translation, degradation, TetR binding, TetR unbinding, and expression leakage (Figure 3 A & B). The deterministic model using identical parameters for all genes based on previous models^23^ (Figure 3C & SI Table 2 & 3) indicated no difference between LITer1.0 versus LITer2.0 systems except when allowing DNA-bound TetR to degrade. A stochastic computational model using the Gillespie algorithm in MATLAB and the Dizzy software package for validation (Supporting Information)^45–47^ matched the deterministic model, showing insignificant changes in basal expression for the selected parameters (Figure 3C).

**Figure 3.**
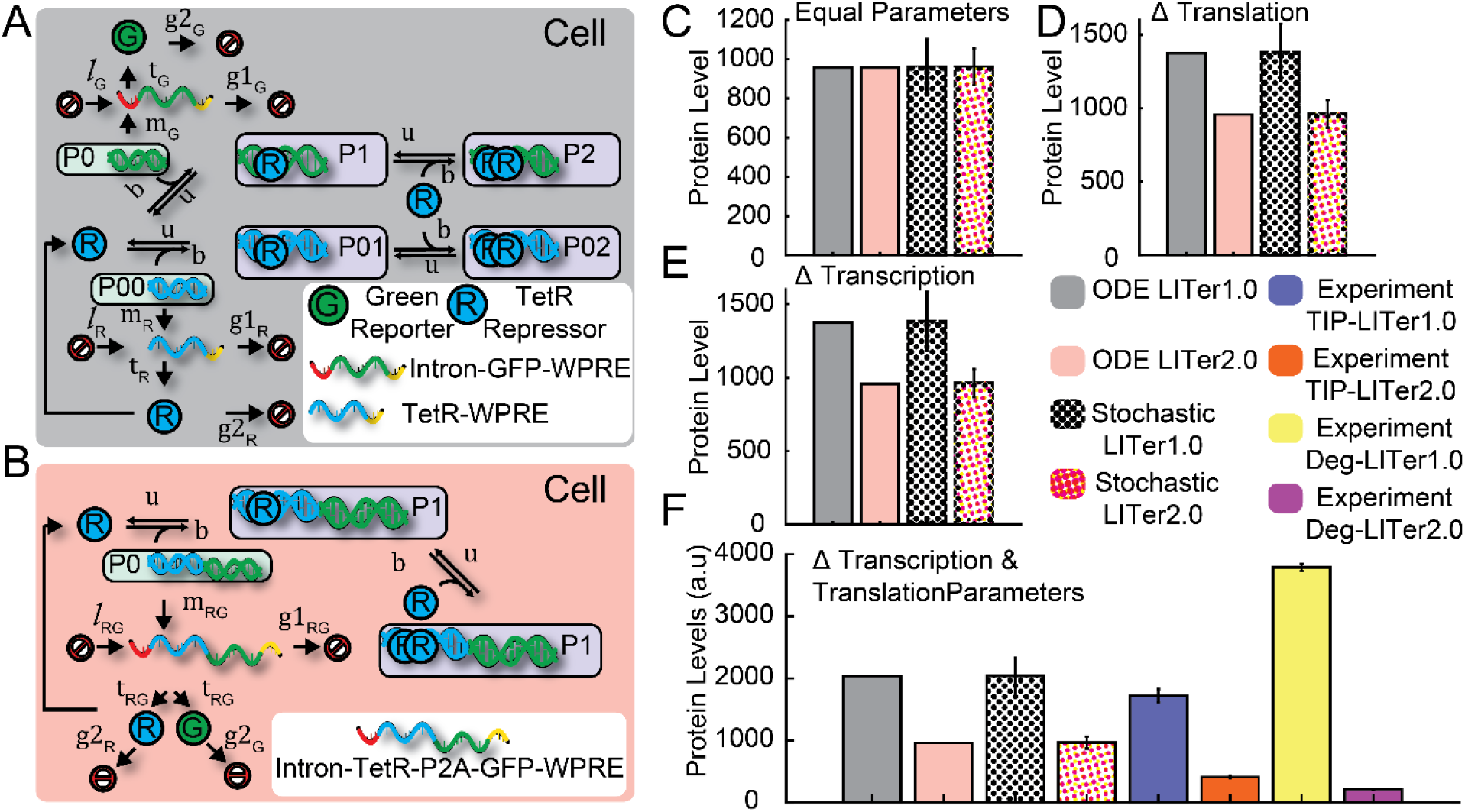
Deterministic & Stochastic Model for exploring parameters to lower basal expression. (A) Schematic illustration of relevant gene circuit reactions used for deterministic and stochastic computational model for two promoter LITer system and (B) one promoter LITer system. The deterministic and stochastic model did not distinguish between degradation or TIP mechanism for controlling TetR. See the Supplement for the notation of rates and chemical species. (C) Bar plot representing deterministic and stochastic model predictions for setting equal parameters between the LITer1.0 & LITer2.0 model for basal expression (no induction). (D) Bar plot representing deterministic and stochastic model predictions for changing the translational parameter between the LITer1.0 & LITer2.0 model for basal expression (no induction). (E) Bar plot representing deterministic and stochastic model predictions for changing the transcriptional parameters (synthesis & leakage) between the LITer1.0 & LITer2.0 model for basal expression (no induction). (F) Bar plot representing deterministic and stochastic model predictions for changing both parameters (transcription & translation) between the LITer1.0 & LITer2.0 model for basal expression (no induction) and accompanying experimental data. 10,000 stochastic simulations were run per condition, and experimental results show 3 replicates for each condition. Bars are means and error bars are standard deviation.

To more realistically match experimental conditions, we then began exploring changing parameters between GFP and TetR production. Despite both GFP and TetR having the same promoters (D2ir), equal parameters in the LITer1.0 models were not realistic because only GFP contains a Kozak sequence and an intron in the 5’ region to increase translational and transcriptional efficiencies respectively, compared to the TetR protein. We therefore moved to explore how such differential transcriptional and translational efficiencies affected the basal expression of GFP in LITer1.0 versus LITer2.0 systems. Thus, to reflect the lack of a TetR intron, we set TetR transcriptional synthesis and leakage rate at 50% of GFP, obtaining a clear drop in basal GFP expression for the LITer1.0 versus the LITer2.0 system (Figure 3D). Next, to reflect the lack of a Kozak sequence, we set TetR translational synthesis rates at 50% of GFP (Figure 3E), which reduced basal GFP expression further in the LITer1.0 versus the LITer2.0 systems.

Finally, we applied both parameter modifications simultaneously to test if greater basal expression reduction occurred (Figure 3F). Indeed, the basal expression decreased more than for each individual parameter change alone. Overall, these models predicted that co-transcribing TetR and GFP would equalize the transcription and translation rates of TetR and GFP, suggesting simple molecular cloning modifications to lower the basal expression of a GOI. Next, we tested these predictions experimentally.

### LILer2.0 gene circuits improve controllability and applicability

To experimentally increase TetR transcriptional and translational efficiency, we incorporated the P2A sequence to allow polycistronic gene expression under a single promoter^27^ (Figure 4A & Supporting Figure S1D & E). We stably integrated the optogenetic gene circuits, and sorted cells into monoclonal cultures termed henceforth TIP-LITer2.0 and Deg-LITer2.0.

**Figure 4.**
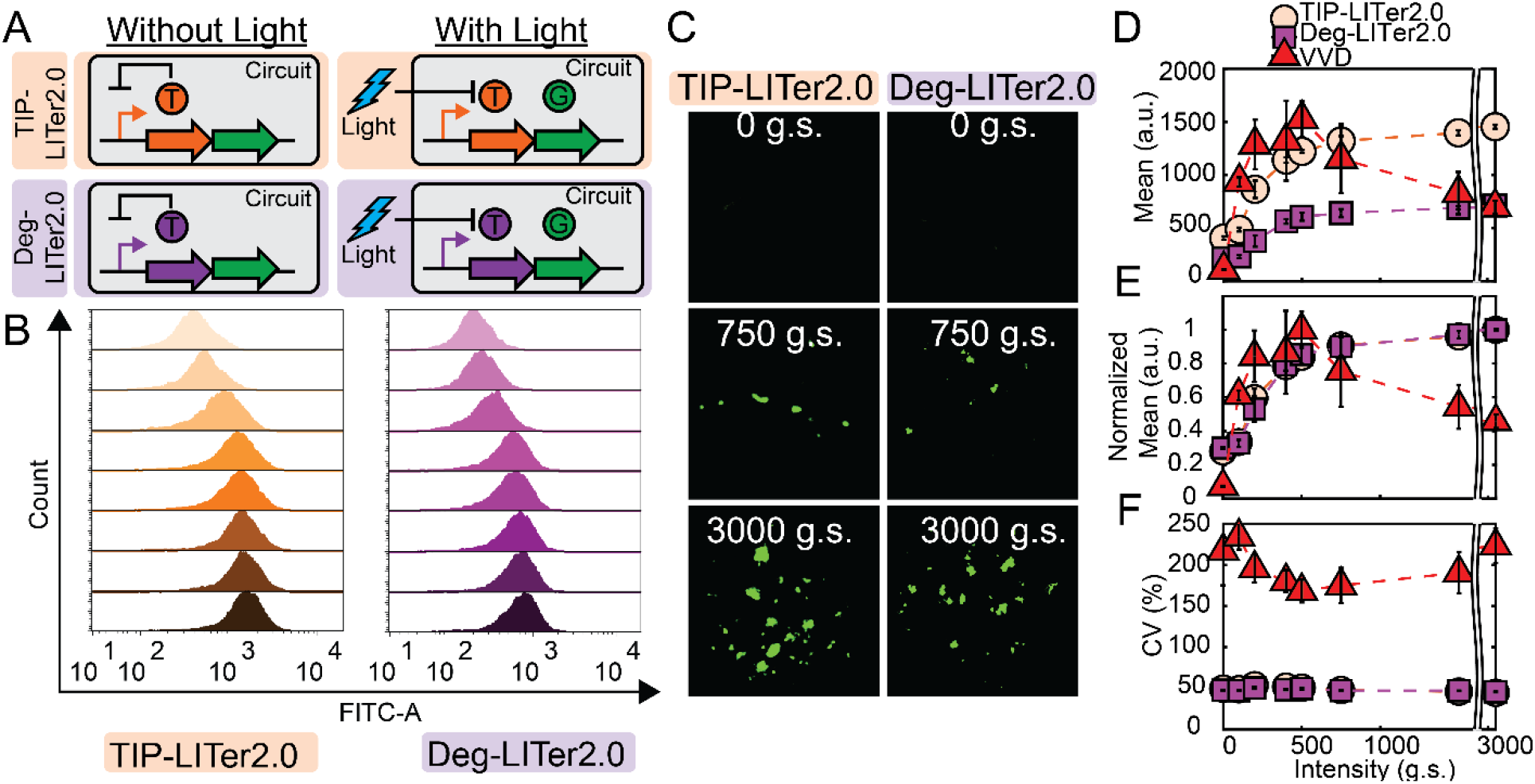
Gene expression control by VVD, TIP-LITer2.0, & Deg-LITer2.0 gene circuits with constant illumination at increasing intensities. (A) Schematic illustration of TIP-LITer1.0 & Deg-LITer1.0 gene circuits stably integrated within Flp-In 293 cell genome. ‘T’ is the TetR regulator with either TIP or Degron and ‘G’ is the Reporter output. (B) Fluorescence histogram distributions from flow cytometry performed on cells with corresponding gene circuit under light intensity titration. (C) Fluorescence microscopy of cells with corresponding gene circuits over light intensity titration. (D) Flow cytometry mean fluorescence expression over light intensity titration for the three gene circuits. (E) Normalized mean fluorescence values to max light intensity level for each gene circuit. (F) Coefficient of variation (CV) over light intensity titration for the three gene circuits. Error bars are standard deviation, N=3. Two-sample t-test was performed on maximum and minimum mean fluorescence data values. TIP-LITer2.0 ON/OFF had a p-value of 2.55E-07 and Deg-LITer2.0 ON/OFF had a p-value of 1.02E-08. Kruskal-Wallis test was performed on CV data set differences. TIP-LITer1.0 vs VVD had a p-value of 2.88E-09 and Deg-LITer1.0 vs VVD had a p-value of 2.87E-09.

To test the performance of these single-promoter LITer2.0 gene circuits, we studied the same illumination regimes as for the LITer1.0 gene circuits, starting with a light intensity titration. In agreement with model predictions, the experimental results indicated that increasing TetR transcription & translation efficiency drastically lowered the basal expression by 82.3% and 97.1% for the TIP-LITer2.0 & Deg-LITer2.0 systems respectively (Figure 3F & 4B). Flow cytometry histograms and fluorescence microscopy indicated tight expression control for the two new gene circuits with over 3-fold dynamic range gene expression induction (Figure 4B-E). Interestingly, the maximum expression flipped compared to the LITer1.0 versions, with the TIP-LITer2.0 having the highest basal expression, while the Deg-LITer2.0 had a similar basal expression to the VVD. Remarkably, despite the massive decrease in basal expression, we found the CV of both the LITer2.0 gene circuits remained nearly identical to the corresponding LITer1.0 versions, remaining up to 5-fold lower than the VVD system (Figure 4F).

To test the effects of light pulsing on the LITer2.0 systems, we next examined how single pulse duration affects the mean and noise of gene expression. The LITer2.0 versions had a similar dynamic range to the LITer1.0 systems (Figure 5A-C). Strikingly, the noise remained up to 5.9 times lower for the LITer2.0 gene circuits compared to VVD (Figure 5D). We also analyzed the effect of the duty cycle (Figure 5E-G) and found around 3-fold dynamic gene expression induction and up to 4.8-fold noise reduction for the LITer2.0 circuits compared to VVD (Figure 5H).

**Figure 5.**
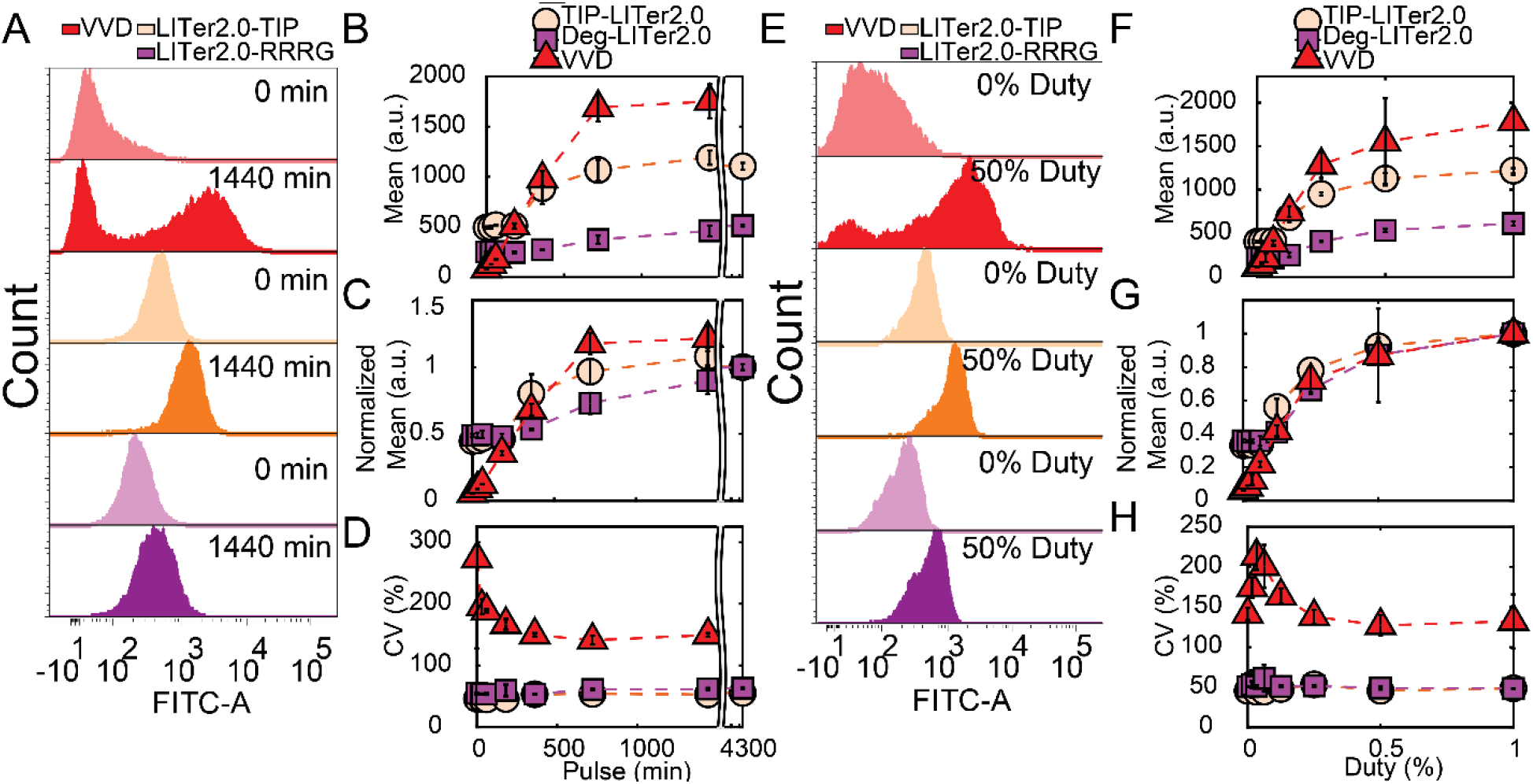
Gene expression control by VVD, TIP-LITer2.0, & Deg-LITer2.0 gene circuits with pulsed illumination. (A) Fluorescence histogram distributions for single light pulse titration. (B) Flow cytometry mean fluorescence expression for single pulse experiment. (C) Normalized mean fluorescence values for single pulse experiment. (D) Coefficient of variation (CV) for single pulse experiment. (E) Fluorescence histogram distributions for duty cycle experiment. (F) Flow cytometry mean fluorescence for duty cycle experiment. (G) Normalized mean fluorescence values for duty cycle experiment (H) CV over duty cycle experiment. Error bars are standard deviation, N=3. Two-sample t-test was performed on maximum and minimum mean fluorescence data values for pulse and duty-cycle experiments. TIP-LITer2.0 ON/OFF had a p-value of 7.72E-06 and Deg-LITer2.0 ON/OFF had a p-value of 3.68E-06 for pulse experiment. TIP-LITer2.0 ON/OFF had a p-value of 1.01E-06 and Deg-LITer1.0 ON/OFF had a p-value of 1.15E-05 for duty-cycle experiment. Kruskal-Wallis test was performed on CV data set differences. TIP-LITer2.0 vs VVD and Deg-LITer2.0 vs VVD had a p-value of 2.88E-09 for the pulse experiment. TIP-LITer2.0 vs VVD had a p-value of 2.88E-09 and Deg-LITer2.0 vs VVD had a p-value of 2.87E-09 for the duty-cycle experiment.

To test whether the four feedback gene circuits still maintained activity to chemical inducers, we also performed dose-response experiments with the small molecule inducer doxycycline (Supporting Figure S7). We found higher fold-induction for all LITer gene circuits (16.8, 4.6, 26.1, and 51.2 for TIP-LITer1.0, Deg-LITer1.0, TIP-LITer2.0, and Deg-LITer2.0 respectively). These results imply that future modifications may further improve light-responsiveness of these gene circuits.

Lastly, as in the LITer1.0 versions, we inspected the linearity of gene expression versus light intensity by fitting a dose-response of a gene circuit to a linear function (Supplement & Supporting Figure S4). The LITer2.0 dose-response was linear up to 225 g.s. from the LPA and improved compared to the LITer1.0 versions. We attribute this improvement to the increase in transcriptional and translational efficiencies for TetR in the LITer2.0 version, therefore matching GFP production. Additionally, Hill functions could fit the full dose-responses of all gene circuits well (Supporting Figure S8), with the Hill coefficient being around 2 for all systems (1.89 [1.46, 2.32], 1.81 [1.5, 2.11], 1.97 [1.62, 2.33], and 2.3 [1.77, 2.83] for the TIP-LITer1.0, Deg-LITer1.0, TIP-LITer2.0, and Deg-LITer2.0 respectively).

Overall, these results show that the LITer2.0 gene circuits are more compact, have lower basal expression, low gene expression noise, and a reasonable dynamic range gene expression induction for various light parameters, making them preferable candidates for precisely controlling expression of a functional gene of interest (GOI).

### Controlling a phenotypically relevant gene with LITer gene circuits

To explore the possibility of using these novel optogenetic tools for expressing a functional GOI, we moved towards adapting the TIP-LITer2.0 system to co-express the mutant oncogene KRAS(G12V) with GFP by incorporating an additional P2A sequence. We called the new gene circuit LITer2.0-KRAS (Figure 6A & Supporting Figure S1F), integrated the system into the Flp-In™ 293 genome, and created monoclonal cells lines.

**Figure 6.**
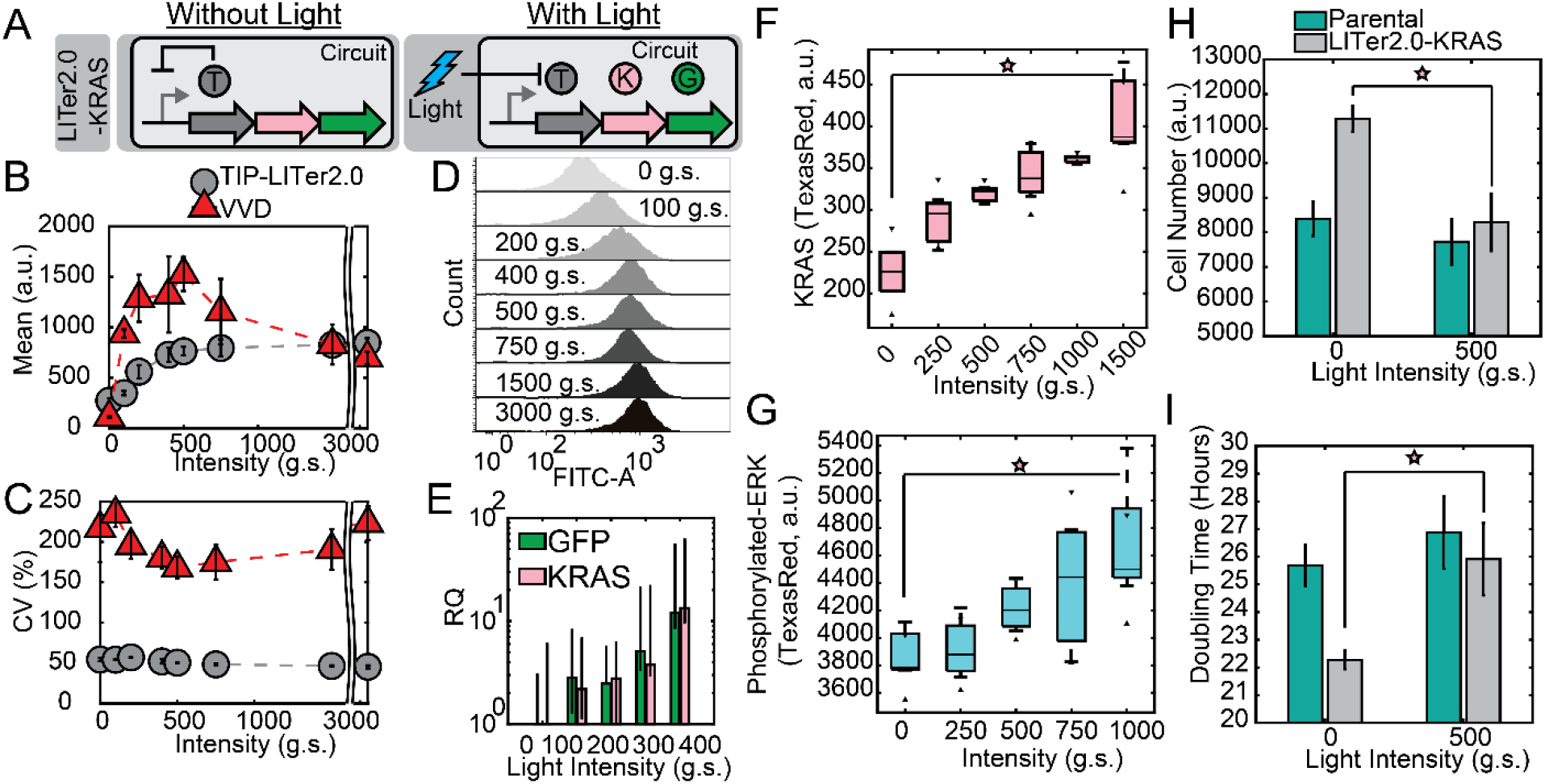
Gene expression control by LITer2.0-KRAS. (A) Schematic illustration of LITer2.0-KRAS(G12V) gene circuit. ‘T’ is the TetR regulator with TIP, ‘K’ is the KRAS(G12V) protein, and ‘G’ is the Reporter output. (B) Mean fluorescence intensity for dose-response. (C) CV for flow cytometry data intensity dose-response. (D) Fluorescence histogram distributions of different light doses. (E) qPCR fold change of KRAS gene circuit induced with varying doses of light expressed on a log(y) axis. (F) KRAS levels measured by flow cytometry with fluorescence-labeled secondary antibody. (G) Phosphorylated-ERK levels measured by flow cytometry with fluorescence-labeled secondary antibody. (H) Growth assay measured by cell number on parental (Flp-In 293) cell line and LITer2.0-KRAS cells 72h after induction. Light causes statistically significant reduction in cell growth compared to basal expression of KRAS cells while light has insignificant effects on the parental cell line. Cells seeded at ~1200 cells per well. (I) Growth assay measured by doubling time on parental (Flp-In 293) cell line and LITer2.0-KRAS cells 72h after induction. Light causes statistically significant increase in cell doubling time compared to basal expression of KRAS cells while light has insignificant effects on the parental cell line. Experiments are performed with three or four technical replicates. Error bars are the standard deviation of replicates. Stars indicate statistical significance. Two-sample t-test was performed on maximum and minimum mean fluorescence data values. LITer2.0-KRAS ON/OFF had a p-value of 3.79E-05. Kruskal-Wallis test was performed on CV data set differences. LITer2.0-KRAS vs VVD had a p-value of 2.88E-09. Two-sample t-test was performed on maximum and minimum KRAS and ERK levels giving a p-value of p-value 0.0071 and 0.014 respectively. Two-sample t-test was performed on cell growth (6H) yielding a p-value of 0.0048.

To validate the gene circuit function, we performed flow cytometry for the LITer2.0-KRAS system which indicated up to 3.1-fold dynamic gene expression induction for GFP and nearly 5-fold noise reduction from the VVD system, (Figure 6B-D). We also analyzed the RNA response of the gene circuit by qRT-PCR, which showed increasing KRAS and GFP expression (Figure 6E) with over an order of magnitude fold-induction.

KRAS is a major activator of growth factor signaling pathways. Usually, post-translational KRAS activity is considered most relevant for downstream signaling, and most studies focus on KRAS-activating mutations. However, increasing KRAS levels should also lead to higher downstream activity, an effect that has been much less studied. The downstream effector ERK is phosphorylated indirectly by KRAS activity through the intermediaries RAF and MEK. If the LITer2.0-KRAS gene circuit produces increasing levels of active KRAS, this should cause rising phosphorylation of ERK in response to light.

To test the direct and downstream effects of KRAS expression tuning, we measured KRAS mean expression levels and the activity of the downstream effector ERK by immunofluorescence. KRAS levels rose nearly 2-fold due to light induction (Figure 6F), indicating that the gene circuit can induce dose-responsive changes in a functional gene and the reporter simultaneously. Likewise, immunofluorescence indicated a rise in the downstream effector phosphorylated-ERK in response to light, confirming that precisely controlled KRAS levels affect downstream signaling (Figure 6G).

Lastly, we checked whether the dose-responsive increases in KRAS levels, and therefore phosphorylated-ERK, could affect cell proliferation as previously demonstrated^48,49^. We grew cells for approximately 72 hours under various light conditions. Afterwards we stained cells with NucBlue and imaged them by microscopy to count their total number per field. Cell growth decreased with increasing light exposure for LITer2.0-KRAS cells, while light did not affect the growth of parental cells significantly (Figure 6H & Supporting Figure S9). Interestingly, we observed that the un-induced basal expression for the LITer2.0-KRAS gene circuit provides some benefit in cell growth compared to parental cells. This indicates that an optimal, low KRAS level may maximize cell growth, while higher KRAS may lead to senescence^50^. Accordingly, converting the cell number into doubling time (Figure 6I), indicated faster growth under low overexpression of KRAS and doubling times reminiscent of parental cells at intermediate doses of light. Lastly, to validate that the observed effects were due to KRAS induction and not light alone, we also analyzed KRAS & phosphorylated-ERK response to doxycycline which recapitulated the light-controlled KRAS findings (Supporting Figure S10).

Overall, the LITer2.0 circuits are beneficial for precision control of intermediate gene expression levels, while maintaining low gene expression noise for a functional GOI, enabling future studies for single-cell control.

## Discussion

Here, we describe five novel optogenetic gene circuits that can be utilized over various light parameters for controlling gene expression levels and reducing noise, enabling single-cell control systems. We report all LITersynthetic gene circuits can drastically reduce noise compared to benchmark tools such as LightOn^28^, facilitating more precise control of gene expression in single cells in the future. Also, we developed six stable monoclonal cells lines that can be utilized for exploring gene expression levels and noise, as well as functional assays involving the RAS pathway in future single-cell investigations.

We show that the systems remain responsive to chemical induction with up to 51-fold change between ON/OFF states, implying that there is ample room to improve circuit architectures for enhanced responses with light. Specifically, we envision these tools can be improved by probing additional peptides with higher affinities, more efficient LOV2 domains, and more efficient degradation tags. We also expect further modeling will lead to architecture manipulations that can achieve a wider dose-response to light, lower basal expression, lower noise, and higher fold-change.

Lastly, the gene circuit components we present here (including small peptides and degradation tags that can be synthesized *in vitro*) offer a platform for modifying existing Tet-based systems into light-responsive tools for controlling single-cells. This will allow gain of spatiotemporal control that has remained cumbersome and restricted by chemical induction alone. Future objectives of this work will aim at scaling the feedback systems to libraries of GOIs and perturbing these GOIs in a spatially restricted manner with single-cell resolution. We also predict that TIP, degradation tags, and other peptides will offer a general-purpose platform that can be utilized for expansion to other common gene circuit architectures including positive regulation, negative regulation, and positive feedback that depend on TetR and rtTA proteins.

## Methods

### Transient transfections

Transient transfections had approximately 75,000 in 500 μL of complete supplemented DMEM media added to each well of a VisiPlate-24 well black plate (PerkinElmer, catalog number: 1450-605) and were first grown for 24 hours. After 24 hours, 500 ng of total plasmid DNA was added to Opti-MEM™ media (Thermo Fisher Scientific, catalog number: 31985062), P3000 reagent, and Lipofectamine 3000 reagent (Thermo Fisher Scientific, catalog number: L3000001), outlined in the Lipofectamine 3000 reagent protocol. Transfection solutions were incubated at room temperature for 10 minutes and added to the cells where mixing occurred by gently shaking. Analysis of all experiments was performed 24-72 hours after transfections were completed.

### Mammalian cell culture

A Flp-In 293 cell line (Thermo Fisher Scientific catalog number R75007) was the parental cell line for all stable-integration cell lines engineered as well as the cell line used for transient transfection. All cell lines were sustained in an environment at 37°C and atmosphere of 5% CO_2_. The cell lines were grown in high glucose Dulbecco’s modified Eagle’s medium (DMEM, Thermo Fisher Scientific, catalog number: 11965-092) that was supplemented with 50mL of 10% Fetal Bovine Serum (FBS, Sigma-Aldrich, catalog number: 12303C), 5 mL of 10,000 units/mL of Penicillin antibiotic and of 10,000 μg/mL of Streptomycin antibiotic (Thermo Fisher Scientific, catalog number: 15140122). Cells with stable gene circuit were maintained under 50 μg/mL of hygromycin drug selection to prevent loss or silencing of integrated payload (Thermo Fisher Scientific, catalog number: 10687010).

### Stable Cell-line Integration

Gene circuits were introduced into Flp-In 293 cells (Thermo Fisher Scientific catalog number R75007) using lipofectamine 3000 (Thermo Fisher Scientific, catalog number: L3000001) according to company protocol with 3×10^5^ cells and 1 μg of plasmid DNA. Cells were transfected with the gene circuit of interest and the Flp recombinase (pOG44, Thermo Fisher Scientific catalog number V600520) plasmid at a ratio of 1:9 respectively. Around 24 hours later cells were washed and given fresh media. Two days later, cells were split to 25% confluency and incubated for several hours, after which hygromycin was added at a concentration of 50 μg/mL. Cells were grown under hygromycin drug selection for several weeks, with media and drug being changed every three to four days. After approximately three weeks, cells were split into T25 filter cap TC flask (USA Scientific, catalog number CC7682-4825) to be frozen down into polyclonal populations or directly sorted into monoclonal populations with flow cytometry. Monoclonal gene circuit populations were expanded and then frozen down according to the Flp-In protocol (Thermo Fisher Scientific).

### Quantitative RT-PCR and quantitative PCR

72 hours post experimental induction, total RNA was extracted from cells from stable-cell lines created using a RNeasy Plus Mini Kit (Qiagen, catalog number: 74134) following the Manufacturer’s protocol. After RNA extraction, reverse transcription was performed using iScript kit (Bio-Rad Laboratories, catalog number: 1708890) following the Manufacturer’s protocol. Following RT-PCR, quantitative PCR was performed using TaqMan Fast Advanced Master Mix (Thermo Fisher Scientific, catalog number: 4444557). TaqMan probes for KRAS and GFP were utilized with FAM dye label (Thermo Fisher Scientific, catalog number: 4331182, Hs00932330_m1 KRAS and Mr04097229_mr EGFP/YFP Assay Id respectively). For normalization, TaqMan probes for glyceraldehyde-3-phosphate dehydrogenase (GAPDH) levels were also utilized with VIC dye label (Thermo Fisher Scientific, catalog number: 4326317E).

### Fluorescence Microscopy

Microscopy was performed 24 or 72 hours after experimental induction. Cell lines were grown on VisiPlate-24 well black plate and imaged on a Nikon Inverted Microscope Eclipse Ti-E and DS-Qi2 camera (14-bit) for phase contrast and fluorescence images. Images were taken with 20x Ph1 objective in phase contrast and GFP mode. The microscope is equipped with Chroma cubes including DAPI 1160B NTE (catalog 49000, Excitation 395/25, Emission 460/50) for DAPI, ET GFP (catalog 49002, Excitation 470/40, Emission 525/50) for FITC/GFP, and ET mCH/TR (catalog 49008, Excitation 560/40, Emission 630/75) for TX Red. Experimental data collection and image analysis was performed using Nikon NIS Elements AR v4.40.00 (Build 1084). All images obtained within a single experiment were collected at the same exposure time per field of interest, underwent identical processing, and normalized to the same fluorescent intensities.

### Flow Cytometry

Prior to flow cytometry, cells were trypsinized with 0.25% trypsin-EDTA (Thermo Fisher Scientific, catalog number: 25053CI) at 37°C for five minutes. After five minutes, trypsin-EDTA was neutralized with supplemented DMEM, after which cells were filtered and read on a BD Fortessa flow cytometer. 10,000 cells per sample were collected per experimental collection.

#### Immunofluorescence

Prior to immunofluorescence staining, cells were trypsinized with 0.25% trypsin-EDTA at 37°C for five minutes. After five minutes, trypsin-EDTA was neutralized with supplemented DMEM. Cells were centrifuged for 5 minutes at 400g. Supernatant was discarded, and cells were resuspended in 1 mL of 4% paraformaldehyde in 1X PBS. Cells were incubated at room temperature for 15 minutes. Cells were then washed with excess PBS, centrifuged for 5 minutes at 400g, and supernatant was then discarded. Cells were then incubated in 1 mL of ice-cold methanol for 30 minutes at −20°C. Cells were then washed with excess PBS, centrifuged for 5 minutes at 400g, and supernatant was then discarded. Cells were then resuspended in 100 μL primary KRAS (Sigma-Aldrich, catalog number: WH0003845M1) or ERK (Cell Signaling Technology, catalog number: 4370S) antibody at a dilution of 1:800 for 1 hour at room temperature in incubation buffer (1X PBS & 0.5g BSA). Cells were then washed with 1 mL incubation buffer PBS, centrifuged for 5 minutes at 400g, and supernatant was then discarded. Cells were then resuspended in 100 μL secondary antibody at a dilution of 1:800 for KRAS (Invitrogen, catalog number: A11005) or 1:2000 for ERK (Cell Signaling Technology, catalog number: 8889S) for 30 minutes at room temperature in incubation buffer. Cells were then washed with 1 mL incubation buffer, centrifuged for 5 minutes at 400g, and the supernatant was then discarded. Cells were resuspended in 500 μL PBS, filtered, and read on a BD Fortessa flow cytometer with10,000 cells per sample.

#### Growth assay

Cells were grown on VisiPlate-24 well black plate at a density of approximately 1200 cells per well. Cells were incubated on an LPA for 72 hours at different light intensities and sustained in an environment at 37°C and atmosphere of 5% CO_2_. After 72 hours, cells were incubated with NucBlue reagent (Thermo Fisher Scientific, catalog number: R37605) and imaged under a Nikon Inverted Microscope Eclipse Ti-E. Bright spot detection was performed on each replicate under DAPI channel excitation, quantified into number of cellular objects, and analyzed for cell number within MATLAB.

#### Statistics and reproducibility

Two-sample t-tests were performed for fold-change calculations between maximum and minimum states of various gene circuits. Additionally, two-sample t-tests were performed for growth assays, KRAS levels, and ERK levels. Kruskal-Wallis test was performed for analyzing significance in CV differences between gene circuits and various light doses.

#### Software

Flow cytometry data was analyzed using FCS Express. Fluorescence microscopy data was analyzed using Nikon-Elements AR package. qRT-PCR data was analyzed using Thermo FIsher Scientific’s Relative Quantification App in the Thermo Fisher Cloud. Experimental data was plotted in MATLAB. Computational models were written in MATLAB & in Dizzy with the parameters given in the supplement. To test the accuracy of the deterministic and stochastic model, we analyzed at steady state and performed 10,000 simulations for each stochastic model parameter.

## Acknowledgements

This work was funded by NIH R35 GM122561. We would like to thank Karl Gerhardt for helping construct the first LPA and Harold Bien, Oleksandra Romanyshyn, Kevin Farquhar, and Yiming Wan for comments and suggestions. We thank Professor Weber for the plasmids gifted for LOV2 protein.

## Author contributions

M.G. and G.B. conceived the project. M.G. performed the experiments. M.G. and G.B. analyzed the data and prepared the manuscript. G.B. supervised the project.

## Notes

The authors declare no competing financial interest.

## Supplementary information accompanies this paper

Requests for materials should be addressed to G. B.

## Data availability

a. Plasmid sequence files can be obtained here: https://gitlab.com/mtguinn/Balazsi-Lab
b. Flow cytometry data and microscopy data can be found here: https://openwetware.org/wiki/CHIP:Data

