## Supporting Information for "Noise-Reducing Negative-Feedback Optogenetic Circuits in Mammalian Cells"

### Supplementary Information

Michael Tyler Guinn[1,2,3] and Gábor Balázsi[1,2,*]

[1]Biomedical Engineering Department, Stony Brook University, Stony Brook, NY 11974 USA

[2]Laufer Center for Physical and Quantitative Biology, Stony Brook University, Stony Brook, NY 11794 USA

[3]Stony Brook Medical Scientist Training Program, 101 Nicolls Road, Stony Brook, NY 11794 USA

**Keywords:** Synthetic biology, optogenetics, light-inducible, noise, feedback

[*]Corresponding author
Gábor Balázsi, Ph.D.
Henry Laufer Associate Professor
Laufer Center for Physical & Quantitative Biology
115C Laufer Center, Z-5252
Stony Brook University
Stony Brook, NY 11794


### Table of Contents

Molecular Cloning: 3
Plasmid Construction: 3
Plasmid Maps (SnapGene ®): 5
Stochastic Modeling: 13
Chemical Reactions: LITer1.0 Systems 13
Chemical Reactions: LITer2.0 Systems 14
Deterministic Models 15
Hill Function Fitting: 17
Linearity Assessment 18
Supporting Figures: 19
Supporting Tables: 29
Supporting References: 33

### Molecular Cloning:

#### Plasmid Construction:

**CMV-TetR-LOV-TIP (pMTG01-Amp-FRT):** The hTetR::NLS:EGFP was digested from pDN-D2ir-TNG4kwh[1] and cloned into pcDNA5/FRT (Thermo Fisher Scientific, V652020) using Spe & Sph restriction enzyme sites. This produced pDN-D2irTN2aG5kwh (pTG01). The hTetR fragment was digested from pDN-D2irTN6kwh[1] and cloned into pTG01 using KasI and NotI restriction enzyme sites. This produced vector pCB-D2irTN5kwh (pTG02). The vector pKH546 (pTG03) was a gift from Professor Wilfried Weber[2]. The hTetR sequence was PCR amplified from pTG02 using primers P1 and P2. The LOV-TIP sequence was PCR amplified from pTG03 using primers P3 and P4. The two fragments were combined in overlap PCR and cloned into pTG01 using BamHI & AgeI restriction enzyme sites. This produced CMV-TetR-LOV-TIP vector.

**CMV-TetR-LOV-RRRG-Degron (pMTG02-Amp-FRT):** The vector pKH546 (pTG03) was a gift from Professor Wilfried Weber[2]. The hTetR sequence was PCR amplified from pTG02 using primers P1 and P2. The LOV-RRRG sequence was PCR amplified from pTG03 using primers P3 and P5. The two fragments were combined in overlap PCR and cloned into pTG01 using BamHI & AgeI restriction enzyme sites. This produced CMV-TetR-LOV-RRRG(Degron) vector.

**TIP-LITer1.0 or CMV-TetOx2-TetR-LOV-TIP---CMV-TetOx2-GFP (pMTG03-Amp-FRT):** The CMV-GFP fragment was digested from pDN-D2irG4kwh[1] and cloned into pTG02 using NotI & KasI restriction enzyme sites. This produced pCB-D2ir-GKwh-D2ir-TN5kwh (pTG04). The TetR-LOV-TIP fragment was digested from pMTG01-Amp-FRT and cloned into pTG04 using MluI & EcoRI restriction enzyme sites. This produced the Dual-NF-TIP or CMV-TetOx2-TetR-LOV-TIP---CMV-TetOx2-GFP vector.

**Deg-LITer1.0 or CMV-TetOx2-TetR-LOV-RRRG---CMV-TetOx2-GFP (pMTG04-Amp-FRT):** The TetR-LOV-RRRG fragment was digested from pMTG01-Amp-FRT and cloned into TG04 using MluI & EcoRI restriction enzyme sites. This produced the Deg-LITer1.0 or CMV-TetOx2-TetR-LOV-RRRG---CMV-TetOx2-GFP vector.

**TIP-LITer2.0 or CMV-TetOx2-TetR-LOV-TIP-P2A-GFP (pMTG05-Amp-FRT):** The hTetR-LOV-TIP sequence was PCR amplified from pMTG03-Amp-FRT using primers P6 and P7. The P2A-GFP-Backbone sequence was PCR amplified from pTG01 using primers P8 and P9. The fragments were combined using NEBuilder HiFi DNA Assembly Reaction Protocol (catalog number: E2621S). This produced the TIP-LITer2.0 or CMV-TetOx2-TetR-LOV-TIP-P2A-GFP.

**Deg-LITer2.0 or CMV-TetOx2-TetR-LOV-RRRG-P2A-GFP (pMTG06-Amp-FRT):** The hTetR-LOV-RRRG sequence was PCR amplified from pMTG04-Amp-FRT using primers P6 and P10. The P2A-GFP-Backbone sequence was PCR amplified from pTG01 using primers P8 and P9. The fragments were combined using NEBuilder HiFi DNA Assembly Reaction Protocol (catalog number: E2621S). This produced the Deg-LITer2.0 or CMV-TetOx2-TetR-LOV-RRRG-P2A-GFP vector.

**VVD system or 5xUAS-mNeonGreen--CMV-Gal4-VVD-p65 (pMTG07-Amp-FRT):** The vector pGAVPO (pTG05) was a gift from Professor Yi Yang[3]. The vector HIV-pBOB-CAG-mNeonGreen-2xNLS (pTG06) was a gift from Gerald Pao[4]. The vector pU5-Gluc (pTG07) was a gift from Professor Yi Yang[3]. The mNeonGreen sequence was PCR amplified from pTG06 using primers P11 and P12 and cloned into pTG07 using HindIII and ApaI restriction enzyme sites. This produced 5xUAS-mNeonGreen-NLSx2 (pTG08) vector. The FRT-Backbone sequence was PCR amplified from pTG01 using primers P13 and P14. The 5xUAS-mNeonGreen-NLSx2 sequence was PCR amplified from pTG08 using primers P15 and P16. The CMV-Gal4-VVD-p65 sequence was PCR amplified from pTG05 using primers P17 and P18. The fragments were combined using NEBuilder HiFi DNA Assembly Reaction Protocol (catalog number: E2621S). This produced the 5xUAS-mNeonGreen--CMV-Gal4-VVD-p65 vector.

**TIP-LITer2.0-KRAS(G12V) or CMV-TetOx2-TetR-LOV-TIP-P2A-KRAS(G12V)-P2A-GFP (pMTG8-Amp-FRT):** The vector pEGFP-C3-KRAS4B-G12V (pTG09) was a gift from Dr. Mark Philips[5]. The KRAS4B (G12V) sequence was PCR amplified from pTG09 using primers P41 and P42. The P2A-GFP sequence was PCR amplified from pMTG05-Amp-FRT using primers P43 and P44. The Backbone-CMV-TetR-LOV-TIP-P2A sequence was PCR amplified from pMTG05-Amp-FRT using primers P45 and P46. The fragments were combined using NEBuilder HiFi DNA Assembly Reaction Protocol (catalog number: E2621S). This produced the CMV-TetOx2-TetR-LOV-TIP-P2A-KRAS(G12V)-P2A-GFP vector (pMTG8-Amp-FRT).

### Plasmid Maps (SnapGene ®):

pMTG01: CMV-TetR-LOV-TIP

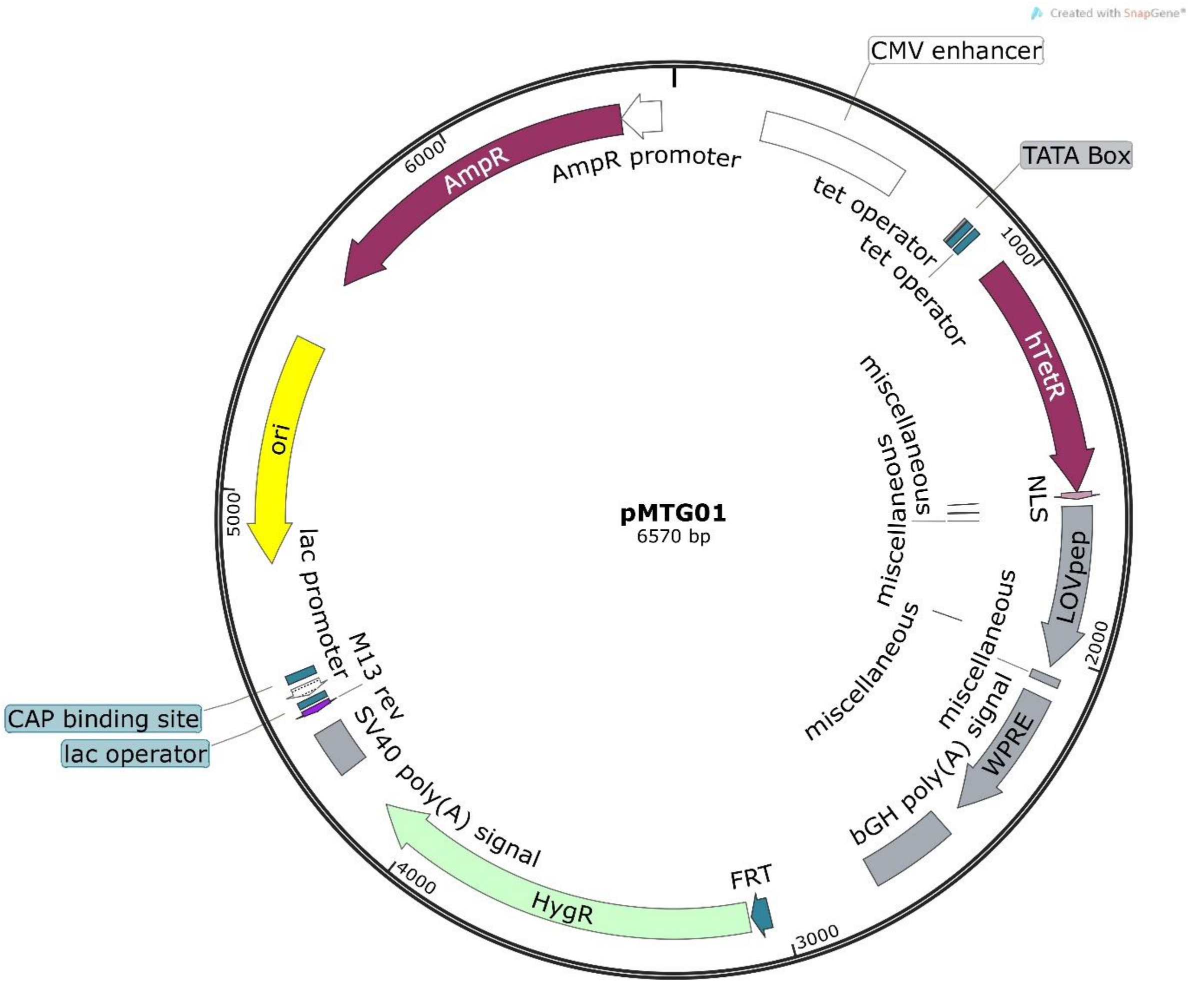

### pMTG02: CMV-TetR-LOV-RRRG-Degron

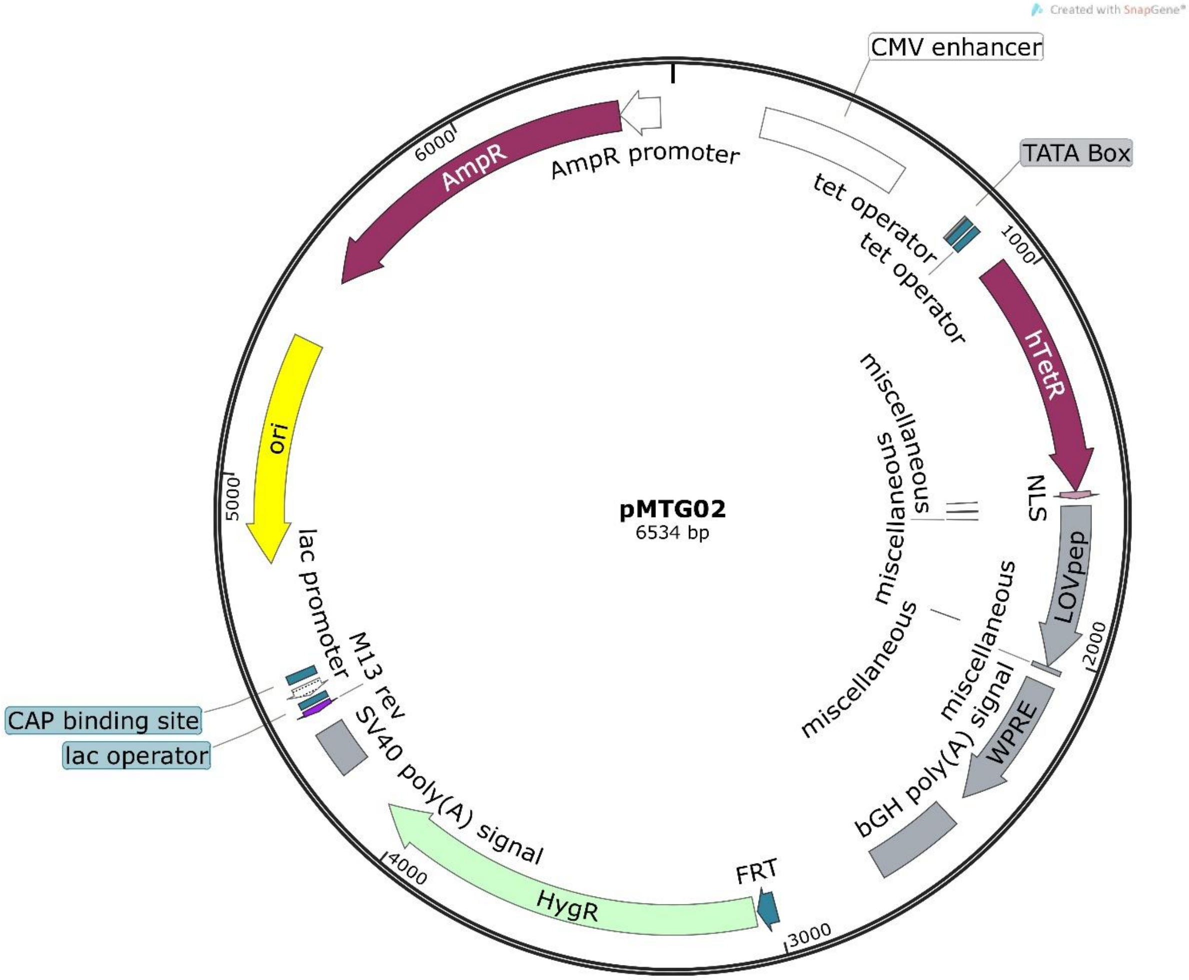

pMTG03: TIP-LITer1.0 or CMV-TetOx2-TetR-LOV-TIP---CMV-TetOx2-GFP

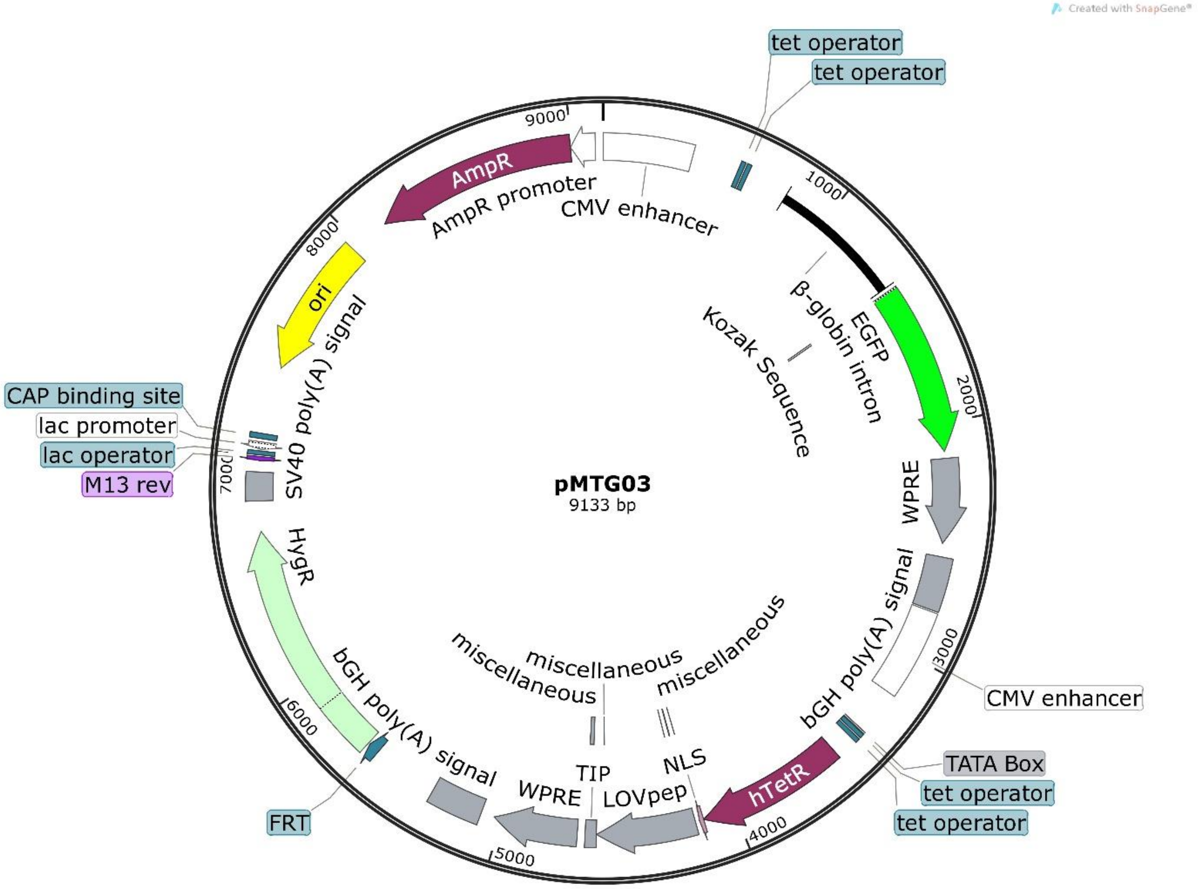

pMTG04: Deg-LITer1.0 or CMV-TetOx2-TetR-LOV-RRRG---CMV-TetOx2-GFP

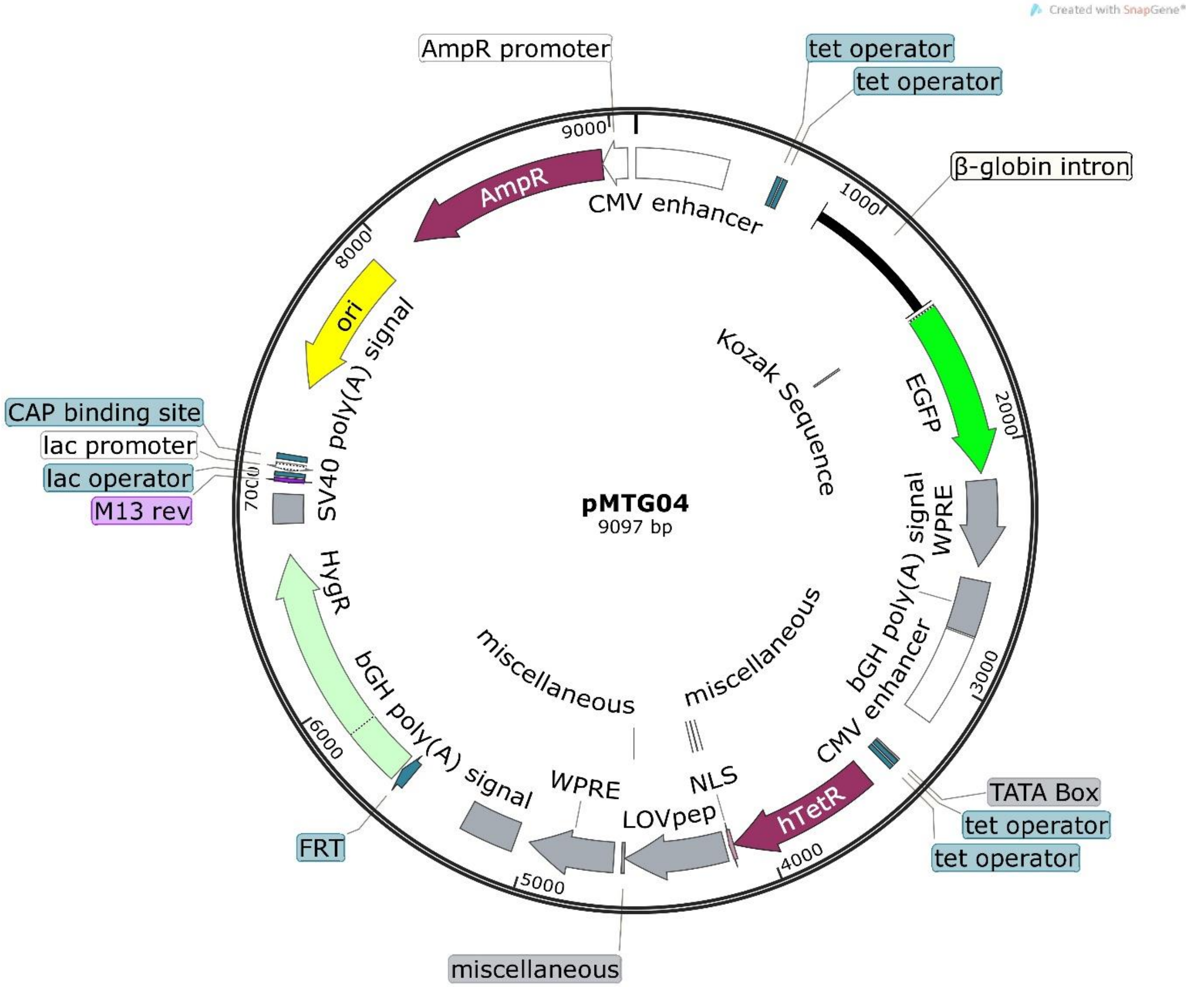

### pMTG05: TIP-LITer2.0 or CMV-TetOx2-TetR-LOV-TIP-P2A-GFP

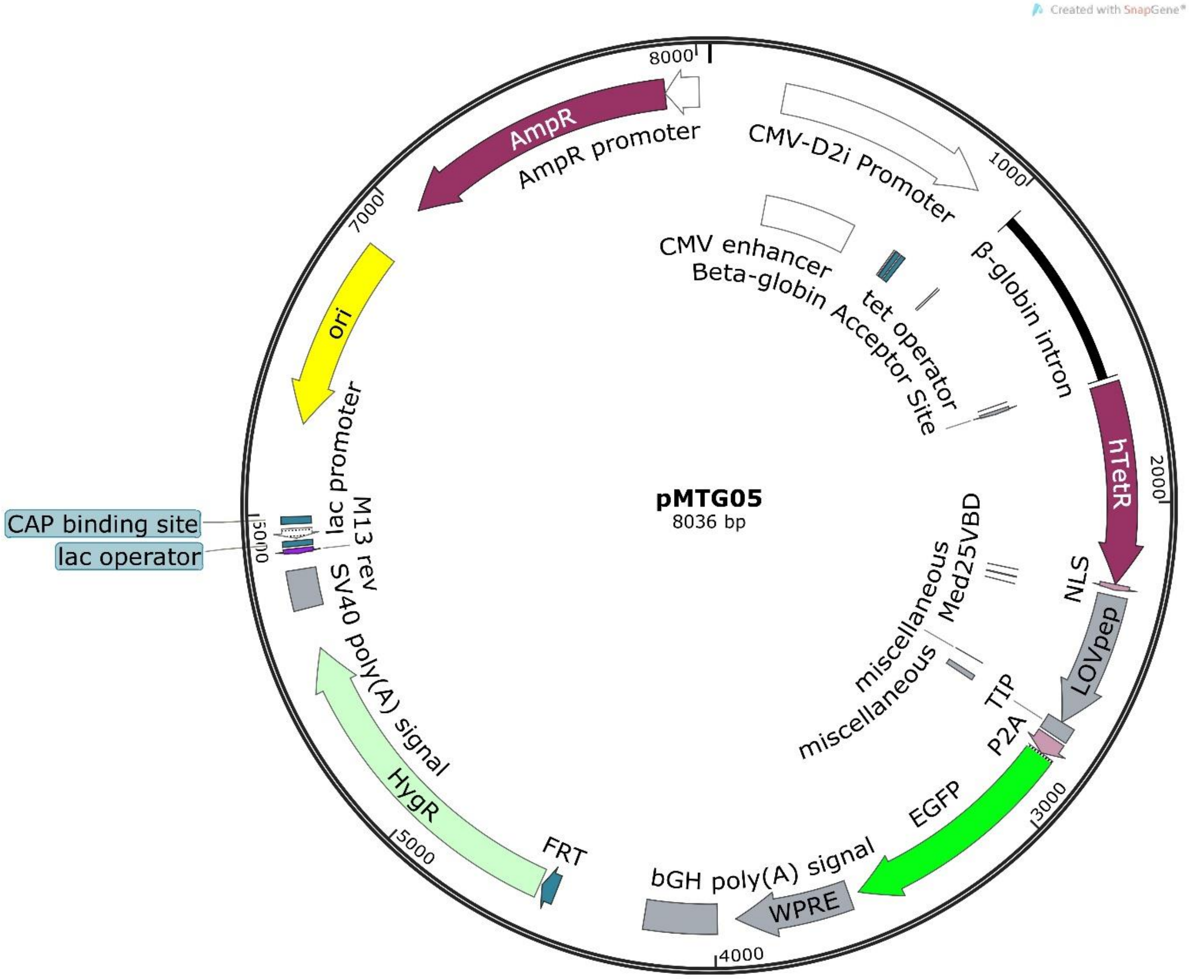

#### pMTG06: Deg-LITer2.0 or CMV-TetOx2-TetR-LOV-RRRG-P2A-GFP

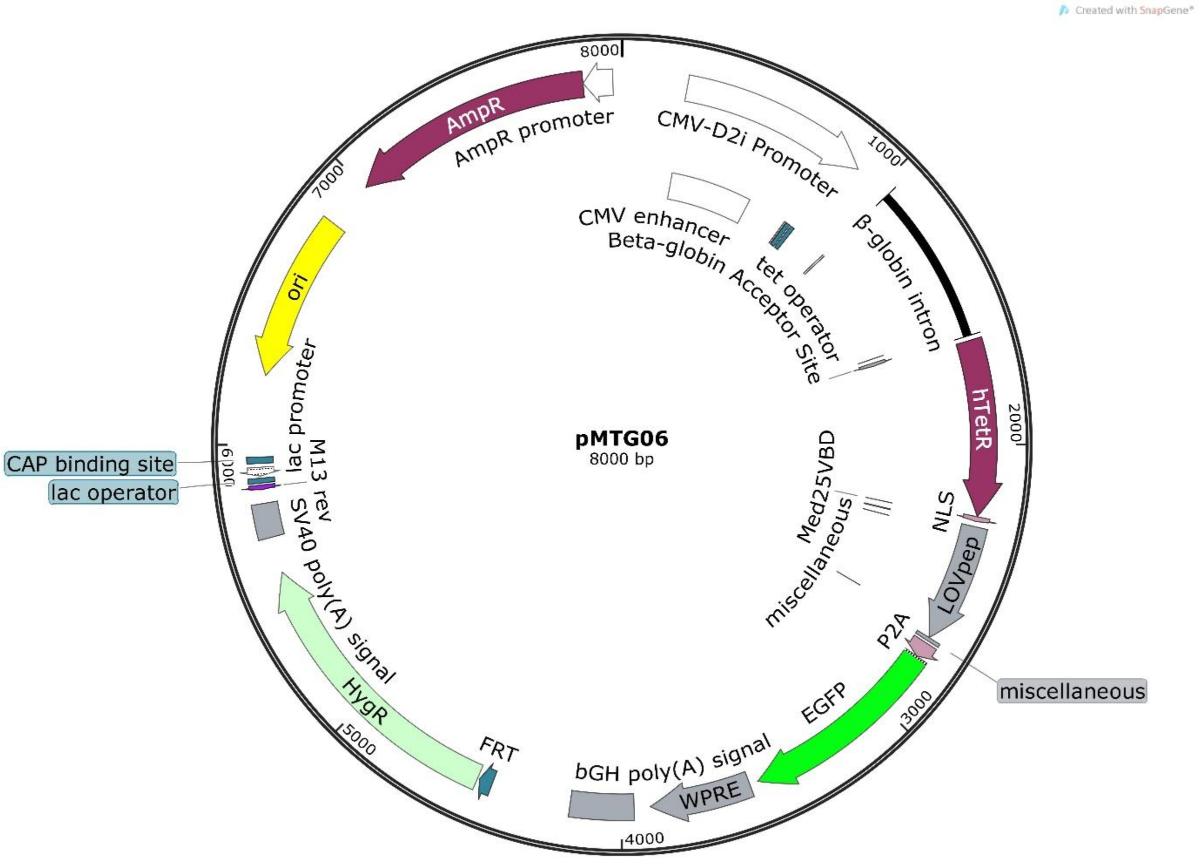

### pMTG07: VVD system or 5xUAS-mNeonGreen--CMV-Gal4-VVD-p65

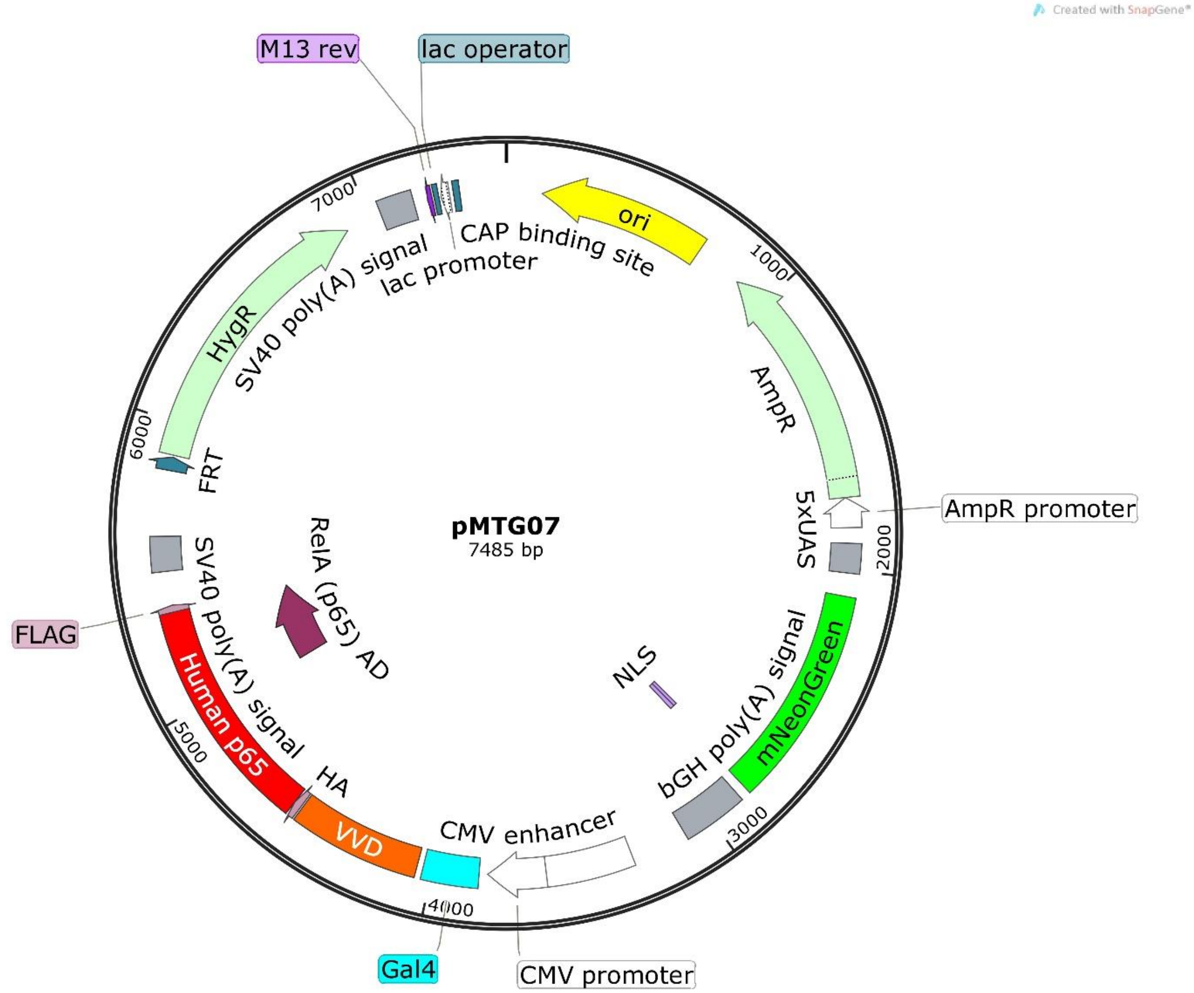

pMTG08: TIP-LITer2.0-KRAS(G12V) or CMV-TetOx2-TetR-LOV-TIP-P2A-KRAS(G12V)-P2A-GFP

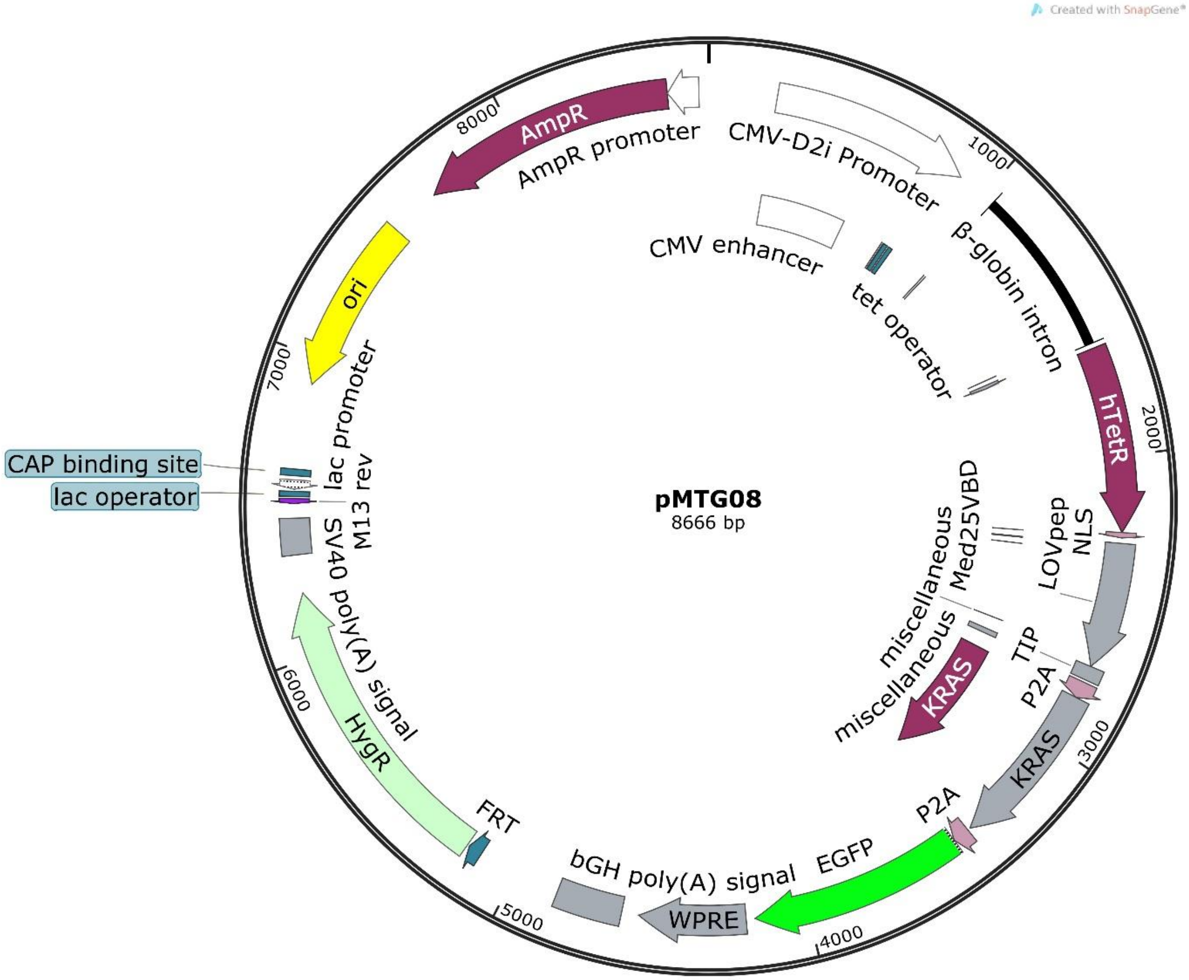

### Stochastic Modeling:

#### Chemical Reactions: LITer1.0 Systems

| | |
|---|---|
| $P00 \xrightarrow{m_R} P00 + M_R$ | Free TetR promoter making TetR mRNA |
| $P0 \xrightarrow{m_G} P0 + M_G$ | Free GFP promoter making GFP mRNA |
| $\emptyset \xrightarrow{l_R} M_R$ | Basal expression of TetR promoter |
| $\emptyset \xrightarrow{l_G} M_G$ | Basal expression of GFP promoter |
| $M_R \xrightarrow{g_1} \emptyset$ | TetR mRNA degraded |
| $M_G \xrightarrow{g_1} \emptyset$ | GFP mRNA degraded |
| $M_R \xrightarrow{t_R} M_R + R$ | TetR mRNA translation |
| $M_G \xrightarrow{t_G} M_G + G$ | GFP mRNA translation |
| $R \xrightarrow{g_2} \emptyset$ | TetR protein degraded |
| $G \xrightarrow{g_2} \emptyset$ | GFP protein degraded |
| $R + P00 \xrightarrow{b} P01$ | TetR binding unbound TetR promoter |
| $P01 \xrightarrow{u} P00 + R$ | TetR unbinding single-bound TetR promoter |
| $R + P01 \xrightarrow{b} P02$ | TetR binding single-bound TetR promoter |
| $P02 \xrightarrow{u} P01 + R$ | TetR unbinding double-bound TetR promoter |
| $R + P0 \xrightarrow{b} P1$ | TetR binding unbound GFP promoter |
| $P1 \xrightarrow{u} P0 + R$ | TetR unbinding single-bound GFP promoter |
| $R + P1 \xrightarrow{b} P2$ | TetR binding single-bound GFP promoter |
| $P2 \xrightarrow{u} P1 + R$ | TetR unbinding double-bound GFP promoter |

Species in **Supporting Table 2**

### Chemical Reactions: LITer2.0 Systems

| | |
|---|---|
| $P0 \xrightarrow{m_{RG}} P0 + M_{RG}$ | Free TetR promoter making TetR/GFP mRNA |
| $\emptyset \xrightarrow{l_{RG}} M_{RG}$ | Basal expression of TetR/GFP promoter |
| $M_{RG} \xrightarrow{g_1} \emptyset$ | TetR/GFP mRNA degradation |
| $M_{RG} \xrightarrow{t_{RG}} M_{RG} + R + G$ | TetR/GFP mRNA translation |
| $R \xrightarrow{g_2} \emptyset$ | TetR protein degradation |
| $G \xrightarrow{g_2} \emptyset$ | GFP protein degradation |
| $R + P0 \xrightarrow{b} P1$ | TetR binding unbound TetR/GFP promoter |
| $P1 \xrightarrow{u} P0 + R$ | TetR unbinding single-bound TetR/GFP promoter |
| $R + P1 \xrightarrow{b} P2$ | TetR binding single-bound TetR/GFP promoter |
| $P2 \xrightarrow{b} P1 + R$ | TetR unbinding double-bound TetR/GFP promoter |

Species in **Supporting Table 2**

### Deterministic Models

The LITer system can be represented by the reaction scheme shown below (using the Degron-based LITer).

Light-activation of the LOV2 domain is at the core of the LITer system. This includes repressor (R)and degron-activated repressor (U). A simplifying step is to assume that light converts the repressor directly into the degron-activted, repressor U.

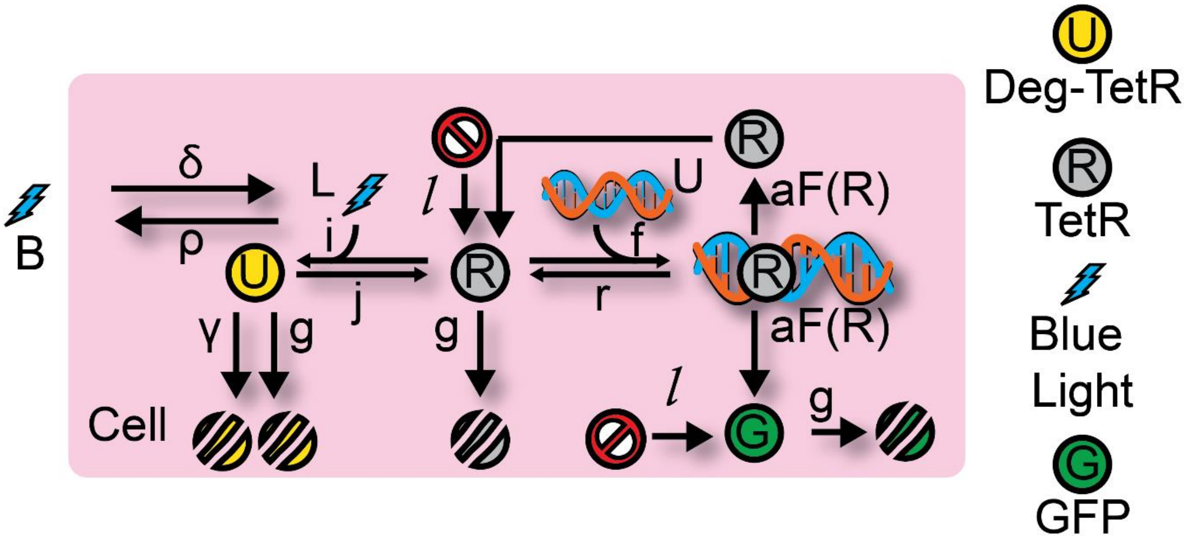


The reactions and ordinary differential equations (ODEs) for this system are:

<u>Symbols:</u>

$R$: $TetR\ repressor$

$U$: $degron\ active\ TetR\ repressor$

$G$: $GFP$

$B$: $Extracellular\ light$

$L$: $Intracellular\ light$

<u>Chemical Equations:</u>

$B \xrightarrow{\delta} L \xrightarrow{\rho} \emptyset$; transfer of light into cell & loss of light

$\emptyset \xrightarrow{aF(R)} R$; synthesis of TetR

$\emptyset \xrightarrow{aF(R)} G$; synthesis of GFP

$R \xrightarrow{iL} U$; LOV2 activation

$U \xrightarrow{j} R$; LOV2 inactivation

$R \xrightarrow{g} \emptyset$; degradation of TetR

$G \xrightarrow{g} \emptyset$; degradation of GFP

$U \xrightarrow{\gamma+g} \emptyset$; degradation of Light active TetR

<u>Differential Equations:</u>

$$\dot{R} = aF(R) - iLR - gR + jU + l$$

$$\dot{U} = iLR - jU - (\gamma + g)U$$

$$\dot{G} = aF(R) - gG + l$$

$$\dot{L} = \delta B - iLR - \rho L$$

This model deviates from the standard Linearizer model in the basal expression term $l$ and the refolding term j, both of which were 0 for the standard model.

The effect of adjusting these parameters away from the standard linearizer can be seen in Supporting Figure 6.

### Hill Function Fitting:

To further inspect gene circuits dose responses, we fit a Hill function to light dose-response data (SE1, Supplement Figure 8).

$$F_x(x) = \frac{ax^b}{c^b + x^b} + d \qquad \text{[SE1]}$$

The ‘a’ parameter represents the theoretical maximum gene expression of the system, the ‘b’ parameter is the Hill coefficient, the ‘c’ parameter is the light value at half-induction, the ‘d’ parameter is the leakiness of the systems, and ‘x’ is the light concentration. We analyzed parameter differences from the Hill function between the different gene circuits (Supporting Table 4). We show the values and confidence intervals as well for all parameters (Supporting Figure 8) which shows all values are statistically significant.

When analyzed from all feedback architectures, we found that the coefficient-squared value around 0.98 or 0.99 for all systems involved. Additionally, when we plotted the 99% confidence interval for the experimental data, fitting the Hill function, we found extremely tight fitting, capturing all data points involved. Overall, these results point towards a characterization of the gene circuit dose-responses and allow end-point users to replace reporters with or add functional GOIs for probing endogenous biological questions.

### Linearity Assessment

Previously, negative feedback and identical promoters were found to produce linearization of the dose response[1,6]. Linearity depended on inducer binding to the repressor, as the main source of decay for repressor and inducer below saturation. This implies that the dilution and degradation of inducer and repressor are relatively small compared to binding of the two into an inducer-repressor complex.

Additionally, non-identical promoters caused deviation of linearity in negative feedback systems[6]. For example, when the promoter driving a reporter gene and a promoter driving a repressor gene differed in the number of Tet operator sites, there was lower linearity between circuit input and output. This contrasted when the promoters and operators were identical, which increased linearity.

In the LITer1.0 system, promoters were identical, however there was an intron and Kozak sequence for the GFP, but not the TetR. This likely altered transcriptional and translational responses to promoter states, causing lower linearity than the original chemical linearizers. Additionally, another source of lower linearity in the LITer systems might be light indirectly affecting TetR through activation of TIP or a degradation tag, while doxycycline directly acts on TetR. There is most likely lower affinity for TIP than the small molecule doxycycline and the light-induced degradation is probably less effective that doxycycline inhibition, lowering linearity from previously observed.

To seek improvement of linearity, we constructed the LITer2.0 systems, where we found higher linearity within both the TIP-LITer2.0 system and Deg-LITer2.0 systems (Supporting Figure S4). Additionally, the most linear range of expression in response to light was between 75 and 225 g.s. on the LPA device shown below.

### Supporting Figures:

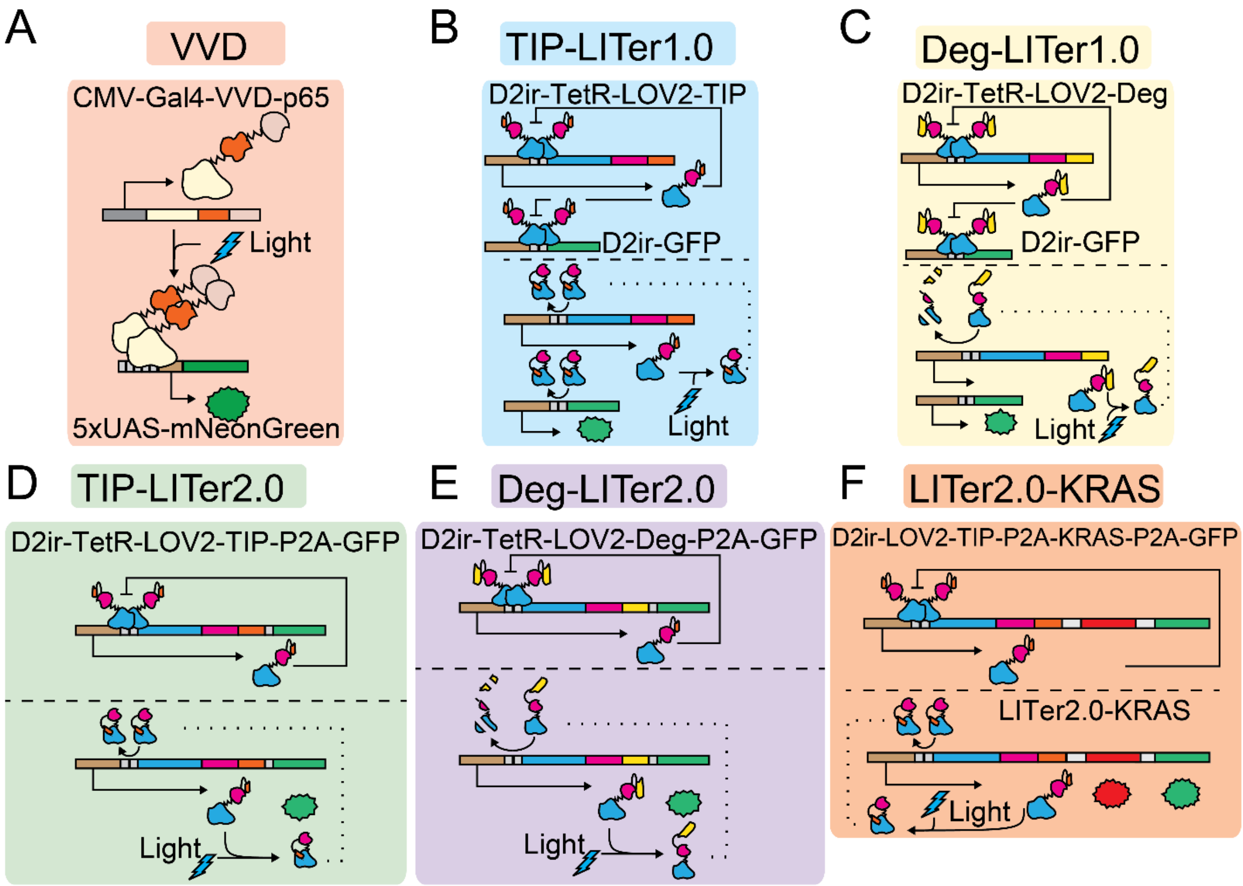


**Supporting Figure S1: Genetic architectures of all six gene circuits.** (A) VVD gene circuit. (B) TIP-LITer1.0 gene circuit. (C) Deg-LITer1.0 gene circuit. (D) TIP-LITer2.0 gene circuit. (E) Deg-LITer2.0 gene circuit. (F) LITer2.0-KRAS

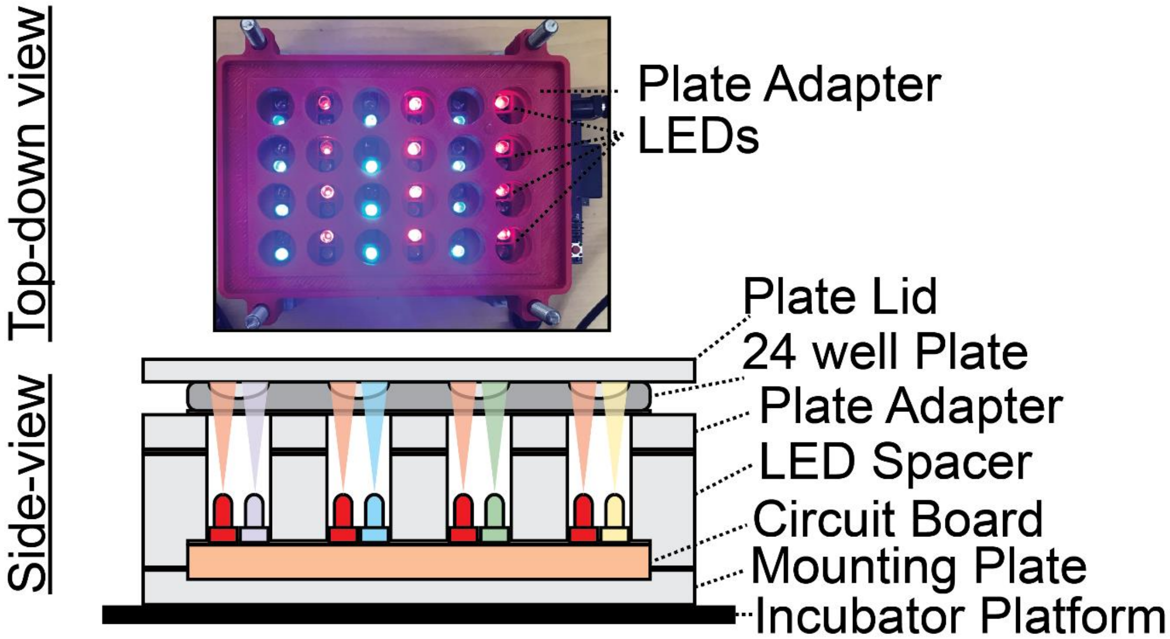


**Supporting Figure S2: Light-Induction Equipment.** The Light Plate Apparatus can contain a mammalian cell culture plate and consists of 24 wells, each of which has two LEDs that can be used for tuning two different wavelengths. Each LED can be turned on continuously, pulsed, or used in complex patterns[7] to induce expression of gene circuit outputs.

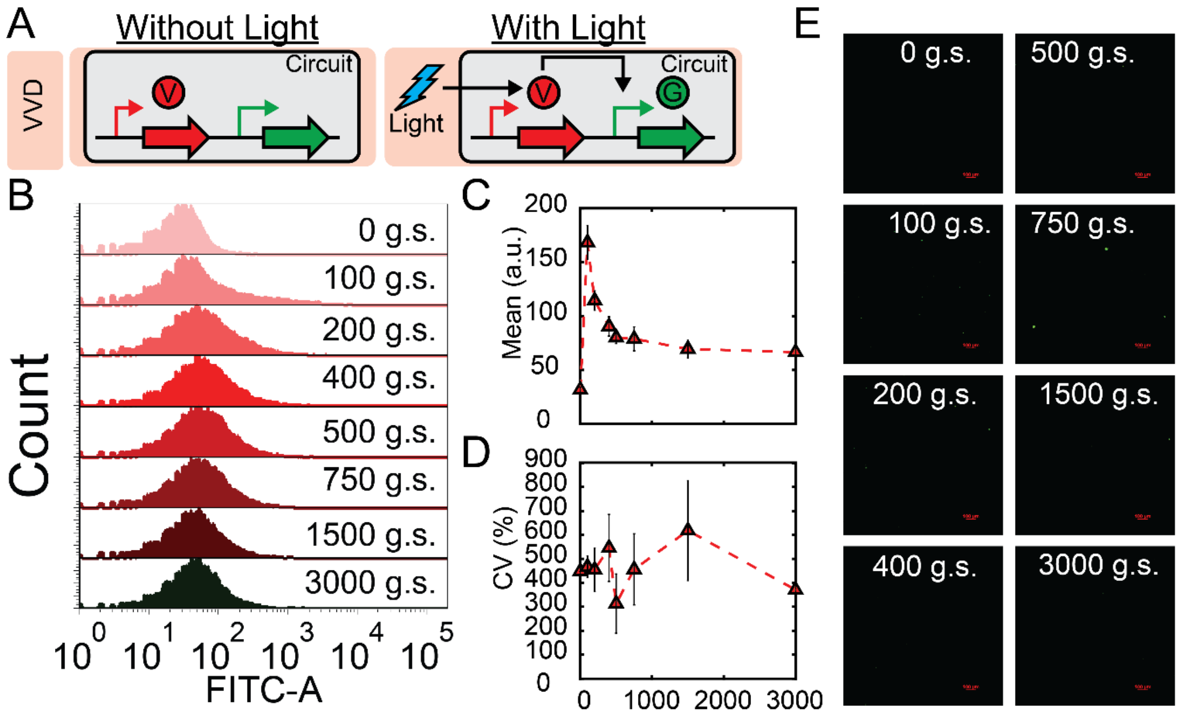


**Supporting Figure S3: Effect of prolonged VVD gene circuit exposure to blue light (72h).** (A) Schematic illustration of VVD gene circuit stably integrated within Flp-In cell genome. (B) Population histogram distributions for flow cytometry performed 72h post light-induction. (C) Flow cytometry mean fluorescence expression over light intensity titration. (D) Coefficient of variation (CV) dose response of gene circuit. (E) Fluorescence microscopy of cells over light intensity titration.

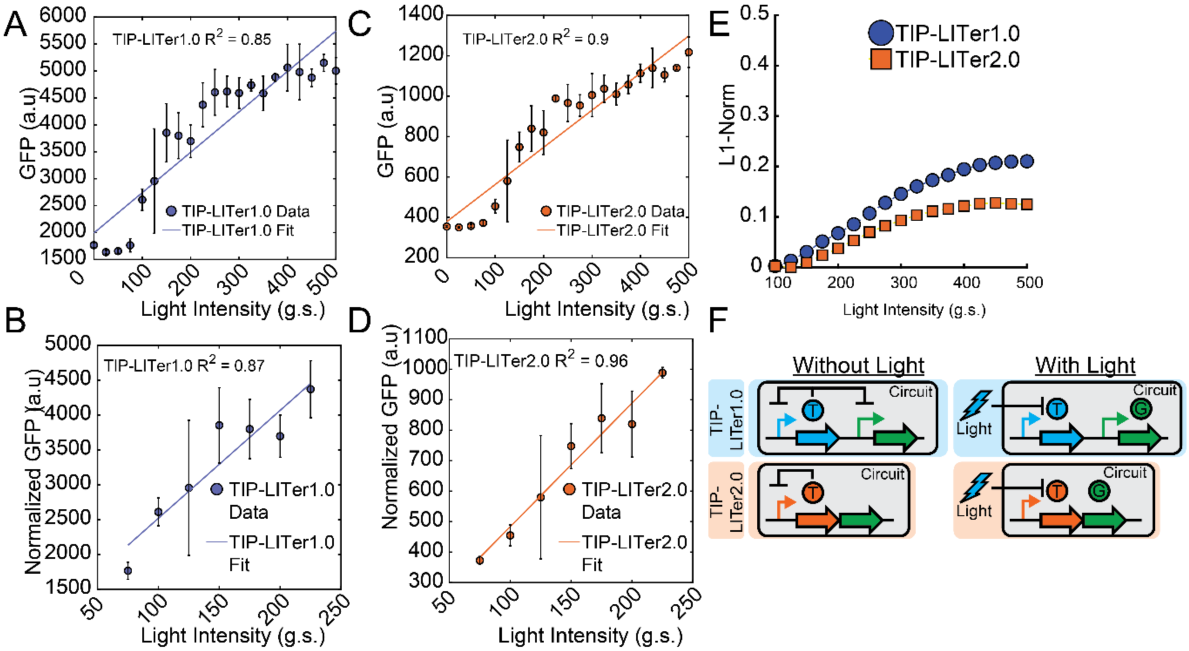


**Supporting Figure S4: Circuit Linearity & Wide-dose response of TIP-LITer1.0 & TIP-LITer2.0. (A)** Wide Intensity dose response assessed for TIP-LITer1.0. (B) Optimal linear range for TIP-LITer1.0 between 75-225 g.s. (C) Wide Intensity dose response assessed for TIP-LITer2.0. (D) Optimal linear range for TIP-LITer2.0 between 75-225 g.s. (E) L1-norm for Intensity dose response for TIP-LITer1.0 & TIP-LITer2.0. (F) Schematic illustration for TIP-LITer1.0 & TIP-LITer2.0.

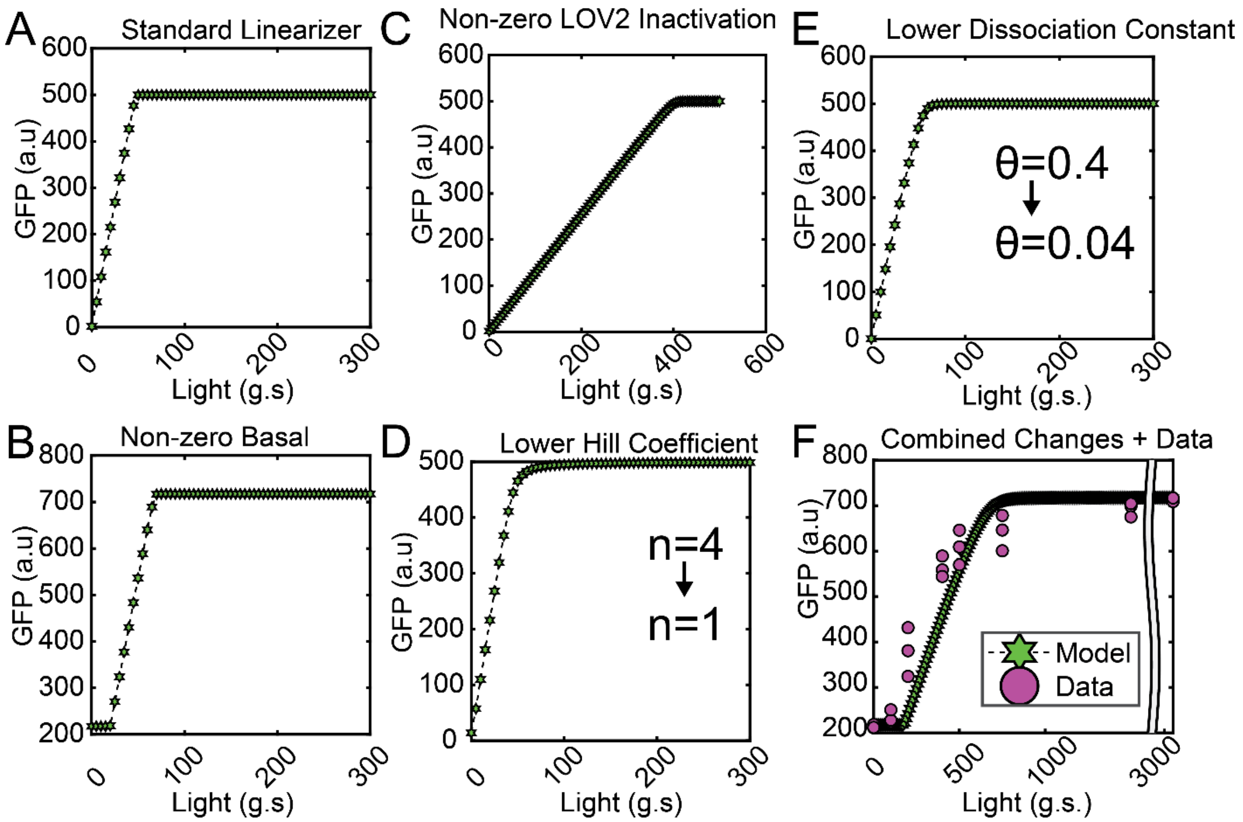


**Supporting Figure S5: Adjustment to Standard Linearizer Model[1].** (A) Standard linearizer model adjusted for light in place of the small chemical doxycycline. (B) Standard linearizer model adjusted to have non-zero basal expression which matches experimental results. (C) Standard linearizer model adjusted to incorporate LOV2 refolding to inactive state, thereby increasing the light needed to induce a level of GFP. (D) Standard linearizer model adjusted to have lower Hill coefficient that results in decreased sharpness of the linear range when approaching saturation. (E) Standard linearizer model adjusted to have lower dissociation constant that results in decreased sharpness of the linear range with approaching saturation. (F) Standard linearizer model adjusted to incorporate all parameters within panels A-E with experimental data from Deg-LITer2.0.

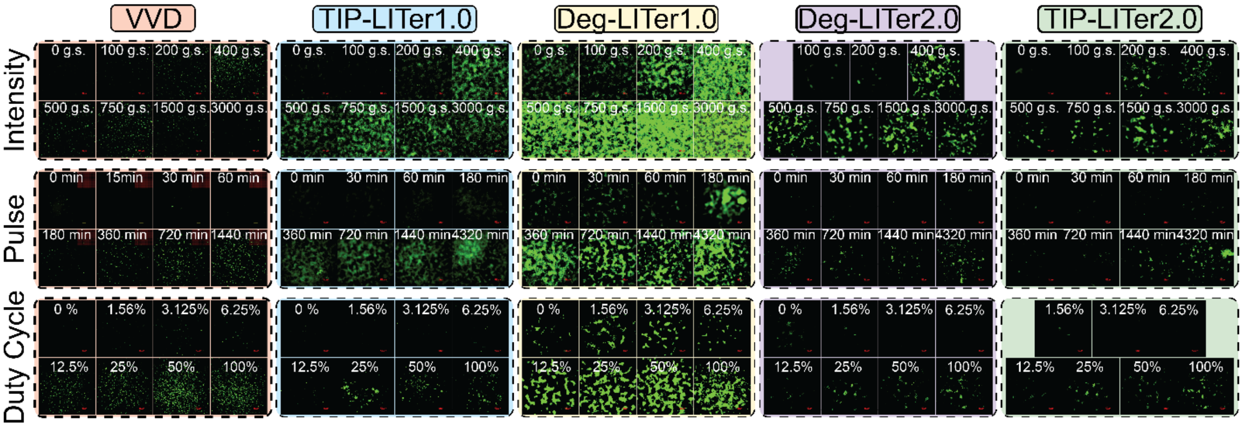


**Supporting Figure S6: Gene circuit dose responses by fluorescence microscopy.** Florescence microscopy from five cell lines integrated with various gene circuits under various experimental conditions including intensity dose response, pulse dose response, and duty cycle dose response. Each grouped column contains a single cell-line of interest with the corresponding gene circuit listed above, while the rows correspond to the light parameter being explored. All cells were exposed to the same exposure time and gain within a given condition (e.g. VVD intensity dose response from 0 g.s. to 3000 g.s.) and normalized accordingly.

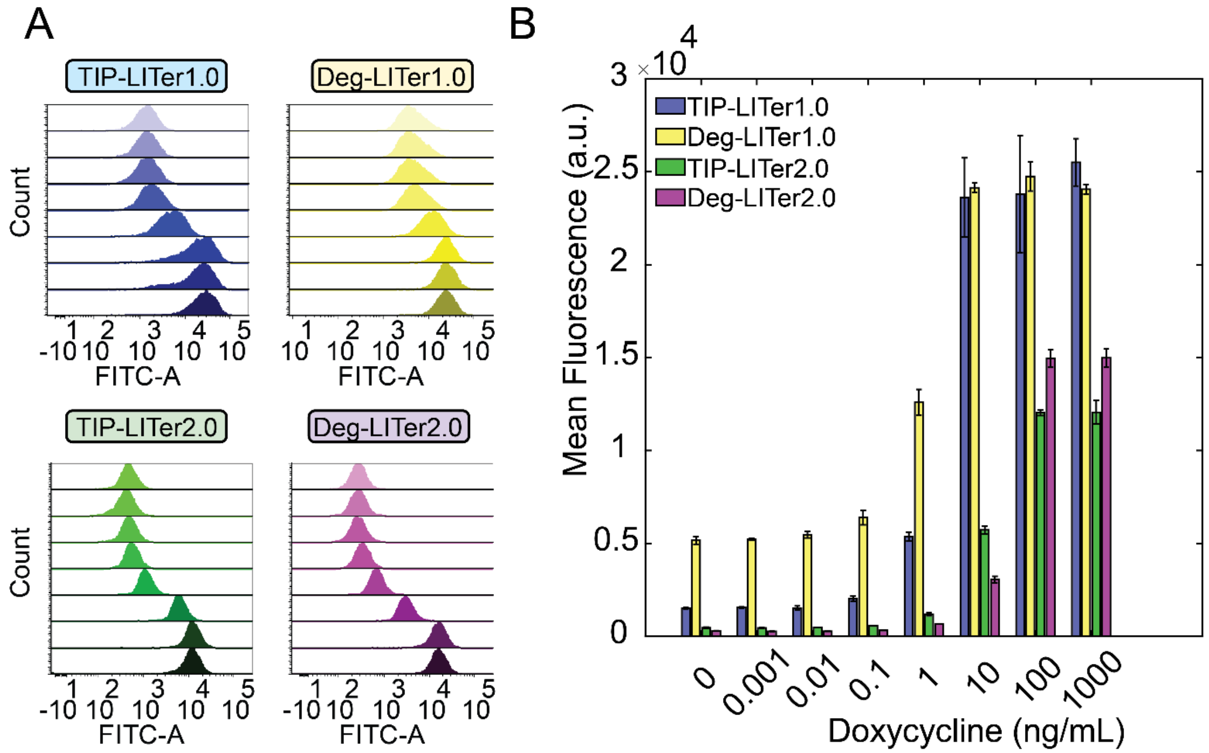


**Supporting Figure S7: Gene circuit dose responses to doxycycline induction.** (A) Flow cytometry histograms showing chemical induction over a wide range of doses, with fold-induction larger than for light other than the Deg-LITer1.0 which showed similar fold-induction for light. (B) Mean fluorescence intensity of the flow cytometry data, showing up to 51-fold change between no induction and max induction.

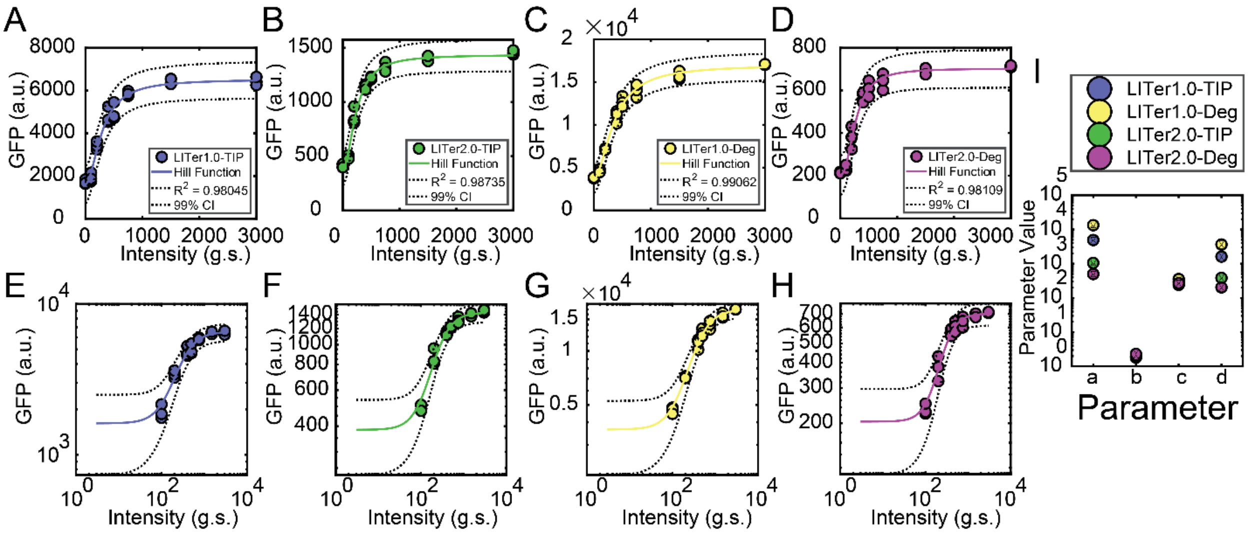


**Supporting Figure S8. Hill function fit to mean expression dose-responses** (A-D) Experimental data for light intensity dose response fitted to Hill Function plotted on linear-linear graph. (E-H) Experimental data for light intensity dose response fitted to Hill Function plotted on log-log graph. (I) Parameters values for Hill function fit for LITer systems shown in Supporting Table 4. Circles represent the mean and black arrows represent the confidence intervals.

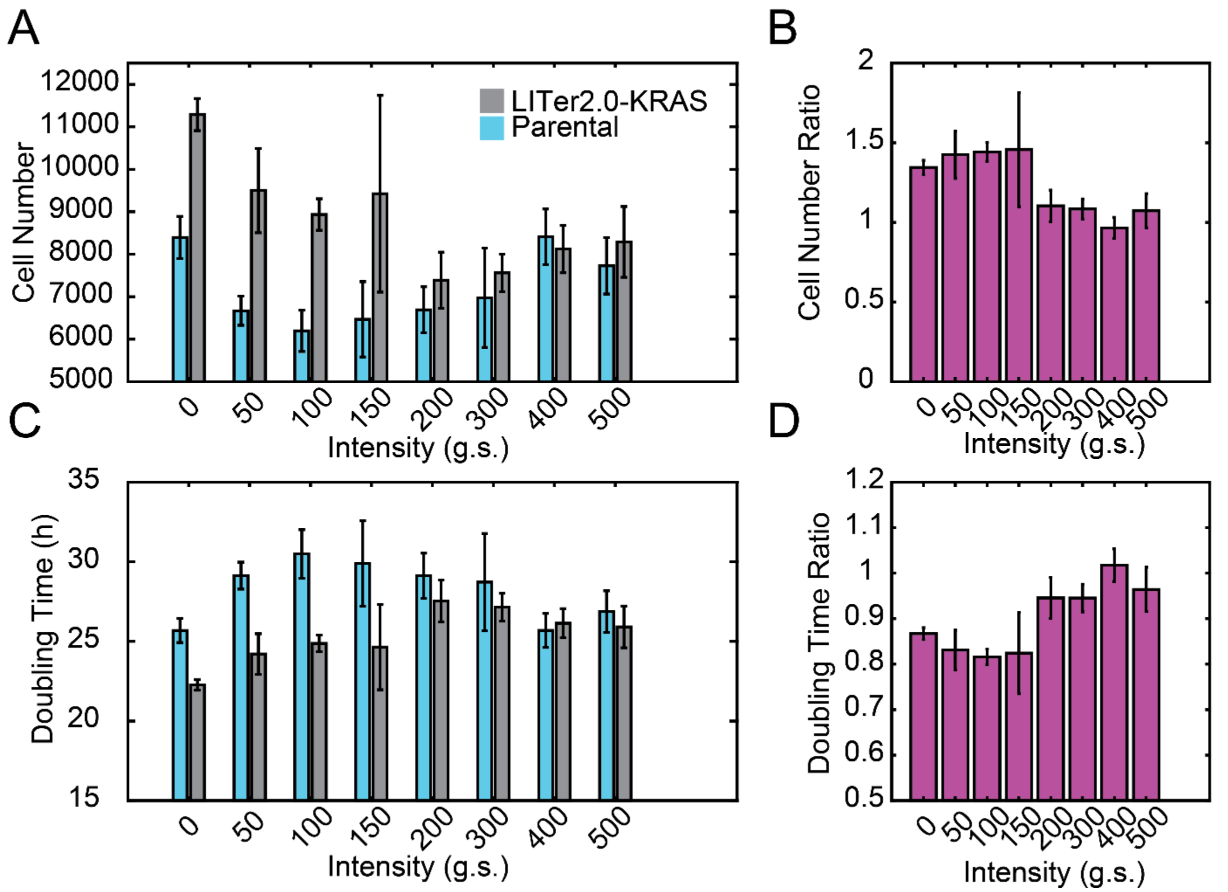


**Supporting Figure S9: Growth assay for LITer-KRAS gene circuit.** (A) Cell number count for parental FLP cells & for LITer2.0-KRAS cells under various light induction. (B) Normalized cell number count for LITer2.0-KRAS cells dived by parental cell count under various light induction. (C) Doubling time for parental FLP cells & LITer2.0-KRAS under various light induction. (D) Normalized doubling time expressed as a ratio of LITer2.0-KRAS cell doubling time divided by parental cell doubling time. Cells were seeded at approximately 1200 cells per well and measured via fluorescence microscopy 72h post-induction by NucBlue staining. Three replicates were performed for each condition.

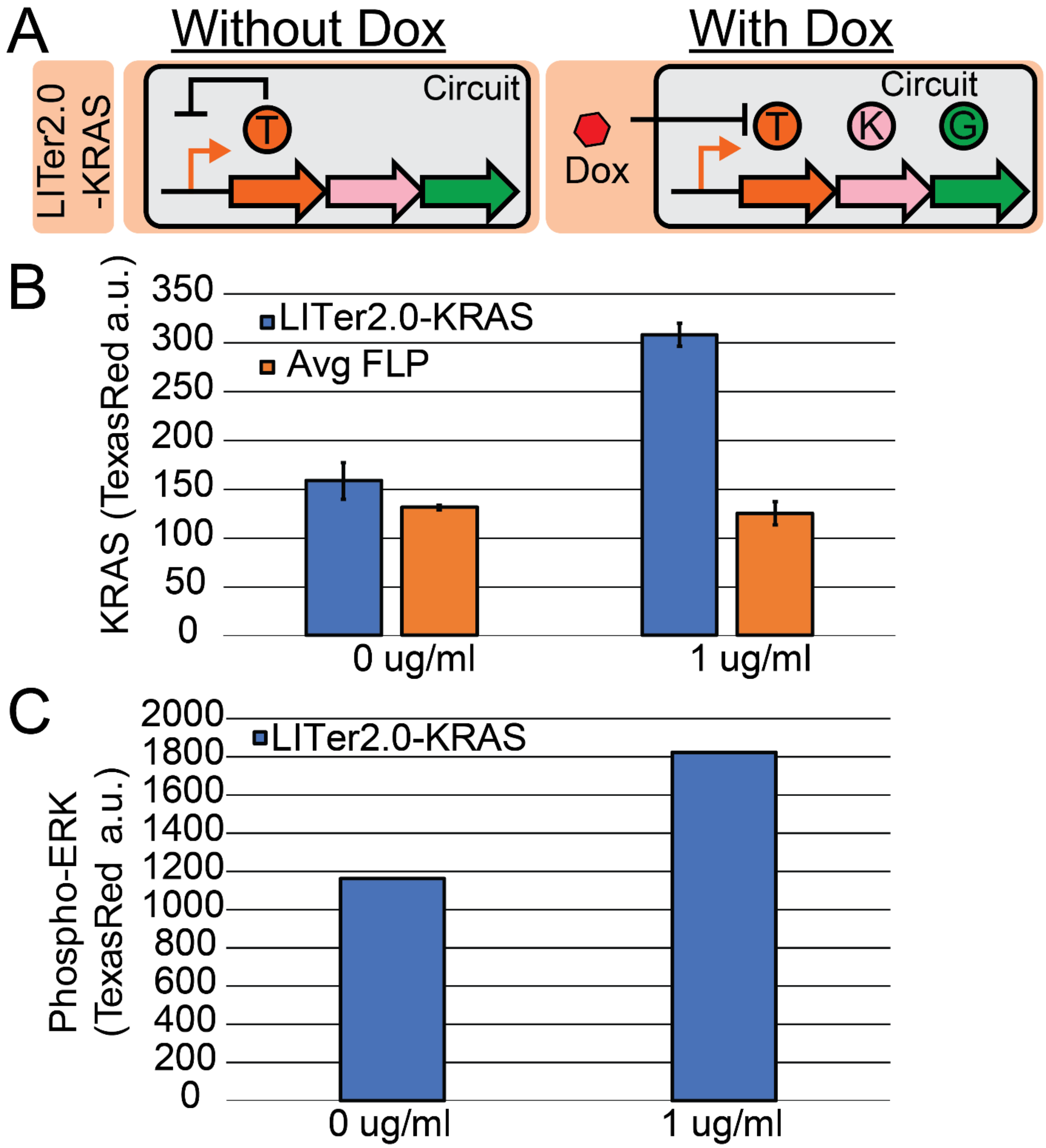


**Supporting Figure S10: Doxycycline induction of KRAS & ERK.** (A) Schematic illustration of LITer2.0-KRAS gene circuit. (B) Doxycycline induction of KRAS in LITer2.0-KRAS cells & parental cell. (C) Doxycycline induction of ERK in LITer2.0-KRAS cells.

### Supporting Tables:

| Primer Name | Sequence | Note |
|---|---|---|
| P1 | CAGTACggatccACCATGGGTTCTAG | BamHI-TetR, forward |
| P2 | gccggcgccgctagcCTTTCTCTTCTTTTTTGGCC | TetR-Lov2 overlap, reverse |
| P3 | GGCCAAAAAAGAAGAGAAAGgctagcggcgccggc | TetR-Lov2 overlap, forward |
| P4 | gtactgACCGGTttaGGAGCCGCCGCCGGAGGGGGCGGCGAAGGCGTAGGCGTTCCAGGTCCAggcggcctcgtcgatg | Lov2-TIP-AgeI, reverse |
| P5 | gtactgACCGGTttagccgcggcggcggg | Lov2-RRRG-degron-AgeI, reverse |
| P6 | caggtcctgcaggcgccaccATGGGTTCTAGACTGGACAAG | hTetR-LOV |
| P7 | tgaagttagtagctccgcttccGGAGCCGCCGCCGGAGG | single promoter TIP |
| P8 | GGTGGCGCCTGCAGGACCTGTAG | pDN-D2irTN2aG5kwh |
| P9 | ggaagcggagctactaacttca | P2A-GFP |
| P10 | TGAAGTTAGTAGCTCCGCTTCCgccgcggcggcgggc | single promoter rrrg |
| P11 | cagtacAAGCTTgccaccATGGTGAGCAAGGGCGAGGA | HindIII-Kozak-tagmNeonGreen, forward |
| P12 | gtactgGGGCCCctaCACCTTCCTCTTCTTCTTAGGCACC | ApaI-NLSx2-tagmNeonGreen, reverse |
| P13 | GTAGCGGCGCATTAAGCGCG | backbone, TG1.55.1 |
| P14 | GACGTCAGGTGGCACTTTTC | backbone, TG1.55.1 |
| P15 | gaaaagtgccacctgacgtcTCGATAGGTACCGAGTTTCTAGACGG | pU5_fragment |
| P16 | aaggccaggaaccgtaaaaaTACCCCCTAGAGCCCCAGCT | pU5_fragment |
| P17 | agctggggctctagggggtaTTTTTACGGTTCCTGGCCTT | pGAVPO_fragment |
| P18 | cgcgcttaatgcgccgctacTTAAGATACATTGATGAGTTTGGACAAAC | pGAVPO_fragment |
| P19 | GAAAAGTGCCACCTGACGTCcaattgcatgaagaatctgcttag | Positive regulation |
| P20 | cgccggcgccgctagcATAAGATCTGAATTCCCGGGATCCGC | Negative regulation (forward use TG1.17.1) |
| P21 | ctaagcagattcttcatgcaattgGACGTCAGGTGGCACTTTTC | Positive regulation, MV |
| P22 | GCGGATCCCGGGAATTCAGATCTTATgctagcggcgccggcg | Negative regulation, MV (reverse use TG1.17.4) |
| P23 | TGGAGGAGAACCCTGGACCTatgcagattttcgtcaagactttgac | fast-GFP |
| P24 | TGATTGCGGCCGCACCGGctatttgtatagttcatccatgcca | fast-GFP |
| P25 | tggcatggatgaactatacaaatagCCGGTGCGGCCGCAATCA | MV |
| P26 | gtcaaagtcttgacgaaaatctgcatAGGTCCAGGGTTCTCCTCCA | MV |
| P27 | GAAAAGTGCCACCTGACGTCcaattgcatgaagaatctgcttag | Positive regulation |
| P28 | cgccggcgccgctagcCCCGGGGAGCATGTCAAG | Positive regulation |
| P29 | CTTGACATGCTCCCCGGGgctagcggcgccggcg | Positive regulation, MV |
| P30 | ctaagcagattcttcatgcaattgGACGTCAGGTGGCACTTTTC | Positive regulation, MV |
| P31 | TTGACCTTGACATGCTCCCCGGGgctagcggcgccggc | LOV-TIP |
| P32 | GCTGAAGTTAGTAGCTCCGCTTCCggagccgccgccgga | LOV-TIP |
| P33 | CCCGGGGAGCATGTCAAGGTCAA | rtTA, PF |
| P34 | GGAAGCGGAGCTACTAACTTCAGC | P2A |
| P35 | GTTCCAGCTGGGCAGCAGGCGGGCCACGGCCATGATGATCTTGCCGGTggcggcctcgtcgatg | LOV-TCP1, use TG1.16.3 forward, PF |
| P36 | GCCTGCTGCCCAGCTGGAACggaagcggagctactaacttcagc | TCP1-P2A, use TG1.16.1 reverse, PF |
| P37 | CTGCTGCCCAGCTGGAACTAActcgagtctagagggccc | MV, PR |
| P38 | CTCTTGTCCAGTCTAGACATggtgaattcggggccgcgga | MV, PR |
| P39 | tccgcggccccgaattcaccATGTCTAGACTGGACAAGAG | rTetR(rtTA), PR |
| P40 | gggccctctagactcgagTTAGTTCCAGCTGGGCAGCAG | rTetR(rtTA)-LOV-TCP1, PR |
| P41 | TGGAGGAGAACCCTGGACCTatgactgaatataaacttgtggtag | KRAS4B |
| P42 | GCTGAAGTTAGTAGCTCCGCTTCCcataattacacactttgtctttgac | KRAS4B |
| P43 | gtcaaagacaaagtgtgtaattatgGGAAGCGGAGCTACTAACTTCAGC | P2A-GFP (Reverse, use TG1.19.4) |
| P44 | GATTGCGGCCGCACCGGTttacttgtacagctcgtccatgcc | P2A-GFP |
| P45 | ggcatggacgagctgtacaagtaaACCGGTGCGGCCGCAATC | MV |
| P46 | ctaccacaagtttatattcagtcatAGGTCCAGGGTTCTCCTCCA | MV (Mother Vector, TIP, use TG1.19.5) |
| P47 | TGGAGGAGAACCCTGGACCTatgatgcaggacgtgtccagctcg | Twist1 |
| P48 | GCTGAAGTTAGTAGCTCCGCTTCCgtgggacgcggacatggacc | Twist1 |
| P49 | ggtccatgtccgcgtcccacGGAAGCGGAGCTACTAACTTCAGC | P2A-GFP |
| P50 | cgagctggacacgtcctgcatcatAGGTCCAGGGTTCTCCTCCA | MV |

**Supporting Table 1. Primers for molecular cloning.**

| LITer1.0 Stochastic Model Parameters | | |
|---|---|---|
| Parameter | Role | Value |
| $l_R$ | TetR mRNA leakage term | $1$ |
| $l_G$ | GFP mRNA leakage term | $2$ |
| $m_R$ | Maximal TetR mRNA production rate | $100$ |
| $m_G$ | Maximal GFP mRNA production rate | $200$ |
| $t_R$ | Maximal TetR protein production rate | $18.75$ |
| $t_G$ | Maximal GFP protein production rate | $37.5$ |
| $g_1$ | mRNA TetR/GFP degradation/dilution rate | $3.5$ |
| $g_1$ | TetR/GFP Protein degradation/dilution rate | $0.046$ |
| $b$ | TetR binding to TetR/GFP Promoter | $0.2$ |
| $u$ | TetR unbinding to TetR Promoter | $20.8$ |
| $M_R$ | Starting # TetR of mRNA | $0$ |
| $M_G$ | Starting # GFP of mRNA | $0$ |
| $R$ | Starting # of TetR Protein | $0$ |
| $G$ | Starting # of GFP Protein | $0$ |

**Supporting Table 2.** Model Parameters

| LITer2.0 Stochastic Model Parameters | | |
|---|---|---|
| **Parameter** | **Role** | **Value** |
| $l_{RG}$ | **mRNA leakage term** | $2$ |
| $m_{RG}$ | **Maximal mRNA production rate** | $200$ |
| $t_{RG}$ | **Maximal protein production rate** | $37.5$ |
| $g_1$ | **mRNA degradation/dilution rate** | $3.5$ |
| $g_2$ | **Protein degradation/dilution rate** | $0.046$ |
| $b$ | **TetR binding to TetR/GFP Promoter** | $0.2$ |
| $u$ | **TetR unbinding to TetR/GFP Promoter** | $20.8$ |
| $M_{RG}$ | **Starting # of mRNA** | $0$ |
| $R$ & $G$ | **Starting # of TetR or GFP Protein** | $0$ |

**Supporting Table 3.**

| Parameters | Mean | 99% CI Min | 99% CI Max |
| --- | --- | --- | --- |
| **a** | 4911.38 | 4443.17 | 5379.58 |
| **b** | 1.89 | 1.46 | 2.32 |
| **c** | 295.11 | 252.55 | 337.67 |
| **d** | 1615.28 | 1305.66 | 1924.89 |
| **a** | 13345.17 | 12409.35 | 14281.00 |
| **b** | 1.81 | 1.50 | 2.11 |
| **c** | 355.24 | 319.86 | 390.62 |
| **d** | 3649.48 | 3088.80 | 4210.15 |
| **a** | 1051.24 | 974.88 | 1127.61 |
| **b** | 1.97 | 1.62 | 2.33 |
| **c** | 240.50 | 212.86 | 268.14 |
| **d** | 383.99 | 329.51 | 438.47 |
| **a** | 499.05 | 454.76 | 543.35 |
| **b** | 2.30 | 1.77 | 2.83 |
| **c** | 269.12 | 234.38 | 303.86 |
| **d** | 203.88 | 172.21 | 235.56 |

**Supporting Table 4.** Hill Function fit parameters for TIP-LITer1.0 (blue), Deg-LITer1.0 (yellow), TIP-LITer2.0 (green), Deg-LITer2.0 (magenta).
